# A High Fraction of Oral Bacteria in the Feces Indicates Gut Microbiota Depletion with Implications for Human Health

**DOI:** 10.1101/2022.10.24.513595

**Authors:** Chen Liao, Thierry Rolling, Ana Djukovic, Teng Fei, Vishwas Mishra, Hongbin Liu, Chloe Lindberg, Lei Dai, Bing Zhai, Jonathan U. Peled, Marcel R.M. van den Brink, Tobias M. Hohl, Joao B. Xavier

## Abstract

The increased relative abundance of oral bacteria detected in fecal samples has been associated with intestinal diseases and digestive disorders. This observation raises two competing hypotheses: either oral bacteria invade the gut bacterial population and expand in the intestine (the *Expansion* hypothesis), or oral bacteria transit through and their relative increase in feces marks a depletion of the gut bacterial population (the *Marker* hypothesis). To address this, we conducted a comprehensive analysis of quantitative microbiome data from mouse experiments and diverse patient cohorts. Our findings consistently support the *Marker* hypothesis as the primary explanation. We further establish a robust inverse correlation between the total fraction of oral bacteria and decreased total bacterial abundance in feces. This correlation underlies the associations between the oral bacterial fraction and multiple patient outcomes consistent with a depleted gut microbiota. By distinguishing between the two hypotheses, our study guides the interpretation of microbiome compositional data and their links with human health.

## Introduction

The microbiomes inhabiting different healthy human body regions have distinct bacterial populations, which reflect the unique characteristics of each ecological niche^1^. Despite occasional exchange among these sites, their resident bacterial populations remain distinctive. For example, humans swallow 1.5 x 10^12^ salivary bacteria per day^2^, some of which can travel to the lower gastrointestinal tract^3–5^ through the enteral^6,7^ and hematogenous^8^ routes. Gastric acids and antimicrobial peptides kill many oral travelers, but those that survive the journey must overcome colonization resistance from gut resident bacteria, which are better competitors in the gut environment^7^. Consequently, oral bacteria are typically scarce in the intestine and hardly detectable in the fecal samples of healthy individuals^9,10^.

Several factors, including antibiotic use, dietary shifts, aging, and intestinal inflammation, can increase the relative abundance of oral bacteria in fecal samples^6^. This relative enrichment of oral bacteria has been associated with a number of diseases, including Crohn’s disease^11^, ulcerative colitis^12^, irritable bowel syndrome^13^, colorectal cancer^14^, and liver cirrhosis^15^. However, the mechanisms underlying these associations remain unclear.

An important point to consider is that the oral bacterial enrichment in the intestine has primarily been detected through amplicon sequencing of the 16S rRNA gene using DNA extracted from fecal samples. This method produces compositional data, and we propose two competing explanations for its interpretation (Fig. 1): either the total abundance of oral bacteria expands within the intestine (the *Expansion* hypothesis), or the absolute abundance of gut resident bacteria decreases, resulting in a relatively higher representation of oral bacteria in fecal samples without a similar increase in their absolute numbers (the *Marker* hypothesis). While the *Expansion* hypothesis suggests that the gut environment has changed to favor oral residents, the *Marker* hypothesis implies that the gut resident population was damaged with reduced total bacterial load. Distinguishing between the two hypotheses is crucial for understanding the microbiome population dynamics and its implications for human health.

**Figure 1:**
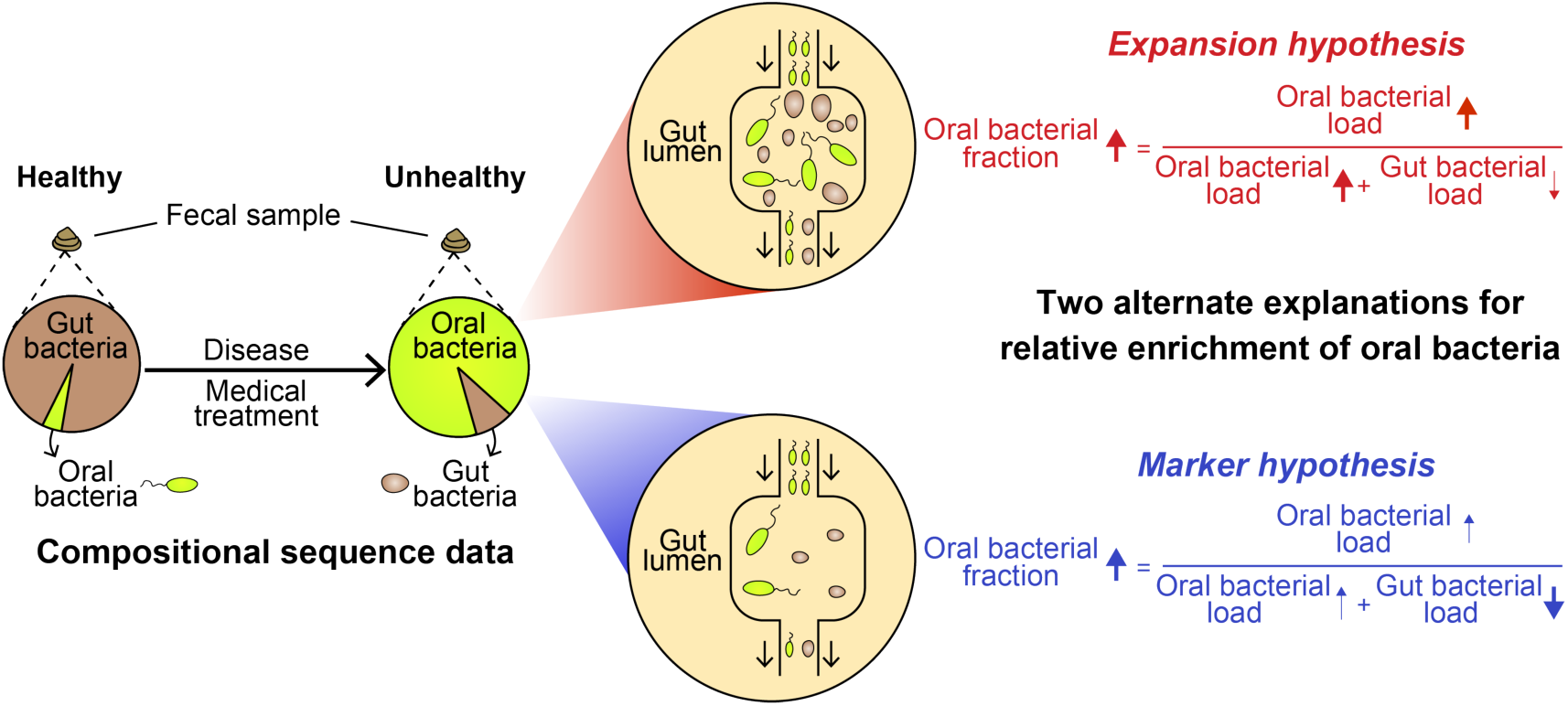
The relative enrichment of oral bacteria in feces has two competing explanations. Certain diseases and medical treatments lead to an enrichment of oral bacteria in fecal samples. This phenomenon has two plausible explanations. In the *Expansion* hypothesis (highlighted in red), the increased fraction of oral bacteria is caused by a similar rise in their absolute numbers. On the other hand, the *Marker* hypothesis (highlighted in blue) attributes this phenomenon to the depletion of gut bacteria. The *Expansion* hypothesis suggests that the gut environment has changed to become more favorable for oral bacteria to establish, whereas the *Marker* hypothesis proposes that perturbations have depleted the gut resident population, and the oral bacteria are simply passing through. Distinguishing between these two hypotheses is crucial for interpreting microbiome population dynamics through compositional analysis of fecal samples.

Here, we compared the two hypotheses using quantitative microbiome data from mice and humans. In the mouse model, we collected paired oral and fecal samples from each subject to quantify the total fraction of oral bacteria detected in the fecal samples after antibiotic treatment or chemically-induced epithelial damage. We find that antibiotic treatment, which depletes the gut bacterial population, increases the fraction of oral bacteria in feces without increasing their absolute numbers. In contrast, chemically-induced inflammation, which alters the composition of the gut microbiota without decreasing its population size, leaves unchanged the fraction of oral bacteria detected in feces. For human patients, we developed a method to estimate the oral bacterial fraction in fecal samples of individuals lacking paired oral samples. Using this method, we observed a similar increase in the relative, but not absolute, abundance of oral bacteria in feces of patients who suffered gut bacteria depletion. These results support the *Marker* hypothesis over the *Expansion* hypothesis. Finally, we show that the fraction of oral bacteria in feces not only serves as a marker for the loss of gut commensal, but is also associated with patient outcomes that involve gut bacterial depletion as a component.

## Results

### Experiments with antibiotic-treated mice support the *Marker* hypothesis

To study the enrichment of oral bacteria in the intestine, we treated eight C57BL/6J female mice with a cocktail of antibiotics (ampicillin, vancomycin, and neomycin) for one week (Fig. 2a). This regimen is known to cause a significant depletion of gut-resident bacteria^16^. We included two additional groups of (a) five untreated mice and (b) five mice treated with DSS (dextran sulfate sodium)—a chemical used to induce epithelial damage without impacting the total bacterial load in feces^17^—for comparison. Throughout the experiment, we collected paired fecal and oral samples, which were subsequently analyzed by 16S amplicon sequencing to profile bacterial composition and by 16S quantitative PCR (qPCR) to estimate total bacterial abundance.

**Figure 2.**
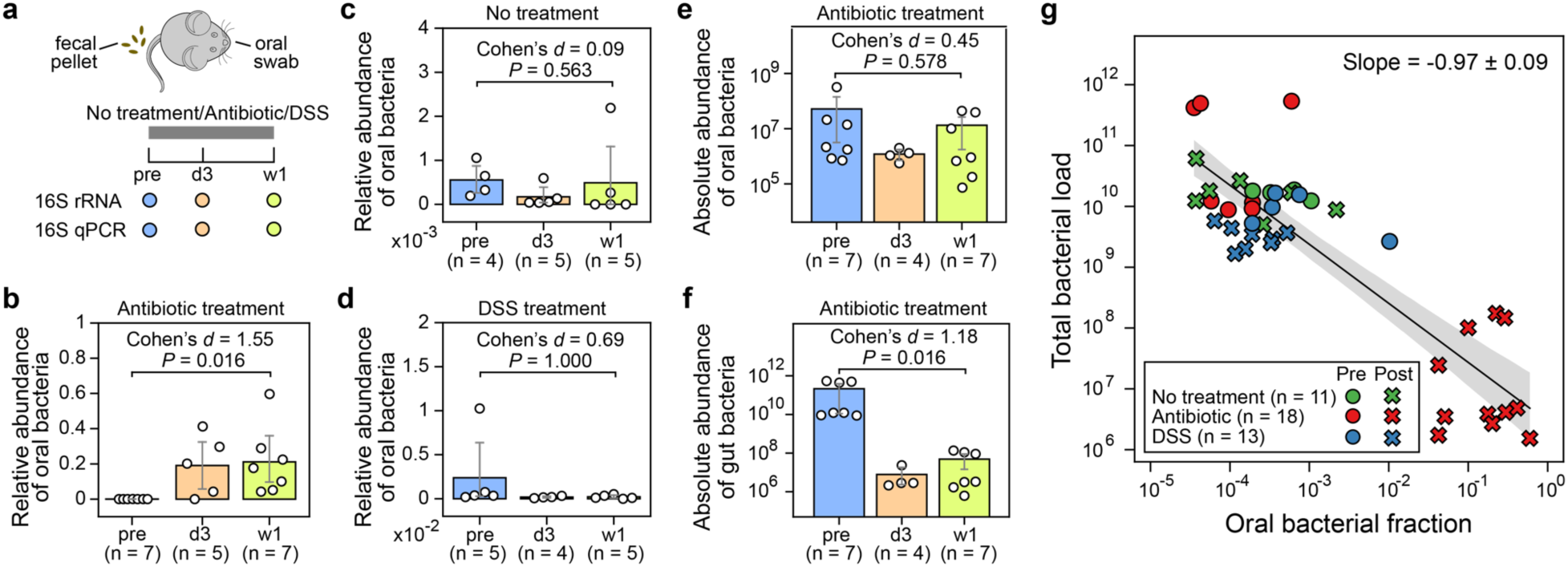
The administration of antibiotics to mice depletes gut bacteria and increases the relative abundance of oral bacteria in fecal samples. **a**, Experimental design. The experiment included three research arms: untreated controls (n=5), mice administered an antibiotic cocktail of ampicillin, vancomycin, and neomycin (n=8), and mice treated with dextran sulfate sodium (DSS) (n=5). Paired fecal and oral samples were collected at three-time points: the same day before treatment initiation (pre), 3 days after treatment initiation (d3), and one week after treatment initiation (w1). Samples with less than 1,000 reads were excluded from analysis (see Extended Data Fig. 1a for excluded samples) and not shown in panels (b)-(g). 16S rRNA gene amplicon sequencing and 16S quantitative PCR (qPCR) data were collected for each sample. **b-d**, Relative abundance of oral bacteria in fecal samples from mice in the antibiotic treatment group (b), no treatment group (c), and DSS treatment group (d). **e**,**f**, Absolute abundance of oral (e) and gut (f) bacteria in fecal samples from antibiotic-treated mice. In panels (b)-(f), each circle represents a fecal sample. Bar heights represent the means, with error bars indicating the 95% confidence interval (CI). *P* values were calculated using a one-sided Wilcoxon signed-rank test. **g**, Linear regression between oral bacterial fraction and total bacterial load in the log_10_-log_10_ scale. Pre-treatment (pre) and post-treatment (d3, w1) samples are represented by circles and crosses, respectively. The black line represents the best linear fit and the shading of the same color indicates the 95% CI. Samples with zero oral bacterial fraction were not shown and excluded from the linear regression analysis. The *y*-axis unit in panels (e)-(g) is 16S copies per gram of feces.

The composition of bacterial populations in the oral and fecal samples before antibiotic treatment were very different, reflecting the distinct habitats of these two organs (Extended Data Fig. 1a). Due to this niche specificity, we could define “oral resident bacteria”, as opposed to “gut resident bacteria”, as those amplicon sequence variants (ASVs) highly abundant and prevalent in pre-treatment oral samples but rarely found in their paired, pre-treatment fecal samples (Methods). We identified 53 such oral ASVs that belong to 32 classified genera, which further enabled us to calculate the total fraction of oral bacteria in all fecal samples (Table S1). We subsequently compared the oral bacterial fraction that was estimated by our approach to the oral bacterial fraction that was calculated by the established source tracking algorithm, FEAST^18^, for post-treatment samples. This comparison yielded a remarkable consistency with an R^2^ value of 0.95 (Extended Data Fig. 2) and validated our approach.

As expected, one week of antibiotic treatment resulted in a significant reduction in the total bacterial load in feces compared to the untreated and DSS-treated mice (Extended Data Fig. 1b). In antibiotic-treated mice, the drop in the total bacterial load coincided with an increased total fraction of oral bacteria detected in the feces (Fig. 2b). This fraction surged from an average of 1.7e-4 in the pre-treatment samples to an average of 0.19 on day 3 and 0.21 after one week. Remarkably, the microbiota composition of post-antibiotic treatment fecal samples was more similar to the pre-treatment oral samples than to the pre-treatment fecal samples (Table S2, Extended Data Fig. 1c). In contrast, there was no enrichment of oral bacteria in the untreated mice (Fig. 2c). In DSS-treated mice, the chemically-induced gut inflammation altered gut microbiota composition (Table S2) but did not deplete the total bacterial load. Consistent with the *Marker* hypothesis, there was no increase in the fraction of oral bacteria detected in feces (Fig. 2d).

We next focused on the antibiotic-treated mice to investigate whether the observed oral bacterial enrichment aligned with the *Expansion* or *Marker* hypothesis. To determine the absolute abundance of oral bacteria, we multiplied the total relative abundance of oral bacteria by the total bacteria load. Notably, the average absolute load of oral bacteria detected in the feces did not increase after antibiotic treatment (Fig. 2e, Extended Data Fig. 3). Instead, the observed relative enrichment of oral bacteria coincided with a significant reduction in the absolute abundance of gut bacteria, which decreased by >1,000 fold on average (Fig. 2f).

The pure *Marker* hypothesis (i.e., oral bacterial load remains constant, and relative enrichment is solely driven by gut bacterial depletion) indicates that the derivative of log_10_(total bacterial load) to the log_10_(oral bacterial fraction) equals -1 (Supplementary Note 1). Conversely, a positive derivative would reject the pure *Marker* hypothesis and favor the pure *Expansion* hypothesis (i.e., gut bacterial load remains constant, and relative enrichment is solely driven by oral bacterial expansion). Remarkably, a linear regression between the log_10_-transformed oral bacterial fraction and total bacterial load in mouse feces produced a slope of -0.97 (Fig. 2g). These results, taken together, strongly support the *Marker* hypothesis as the predominant explanation for the enrichment of oral bacteria after antibiotic treatment.

### Method for quantifying oral bacteria in fecal samples from human subjects without paired oral samples

We next aimed to determine whether the *Marker* hypothesis remains the major mechanism for the reported enrichment of oral bacteria in the human intestine. Similar to the mouse microbiome, bacterial populations residing in the human body have distinct compositions depending on the body site^1^. We, therefore, adopted a similar strategy as used in our mouse data analysis to identify and quantify oral bacteria in human feces (Methods). We analyzed 2,932 paired oral (collected from several oral cavity sites) and fecal samples from 223 healthy individuals in the Human Microbiome Project (HMP)^1^. We found 178 oral ASVs that belong to 42 classified genera, including 24 *Prevotella* and 15 *Streptococcus* ASVs (Extended Data Fig. 4, Table S3). The set of 178 reference oral ASVs obtained from the HMP cohort enabled us to calculate the total fraction of oral bacteria in new fecal samples, even without paired oral samples. We validated our approach by analyzing a publicly available dataset consisting of paired fecal and saliva samples from patients with inflammatory bowel disease and their healthy controls^19^ (Extended Data Fig. 5, Supplementary Note 2).

### Longitudinal analysis of patients hospitalized for allo-HCT supports the *Marker* hypothesis

Having established a method for detecting oral ASVs in human samples, we leveraged a previously compiled human microbiome dataset^20^ to test the two hypotheses. This dataset comprises 10,433 longitudinal fecal samples collected from 1,276 patients who underwent allogeneic hematopoietic cell transplantation (allo-HCT) over the past decade at Memorial Sloan Kettering Cancer Center (MSKCC). Within this dataset, a nested subset of 3,108 samples included 16S qPCR data. Since paired oral samples were unavailable, we used the above reference set to identify oral ASVs and quantify their total fraction in fecal samples (Table S4). Interestingly, 901 of the 10,433 fecal samples were dominated by a single oral ASV with a relative abundance that exceeded 30% (Fig. 3a). These oral ASVs with > 30% abundance primarily belong to three genera, *Streptococcus*, *Actinomyces*, and *Abiotrophia*, which dominated 778, 73, and 30 samples respectively. Notably, factors such as biofilm-forming capacity (Supplementary Note 3) and sequencing depth (Fig. S1) did not account for the observed variations in oral bacterial fraction.

**Figure 3:**
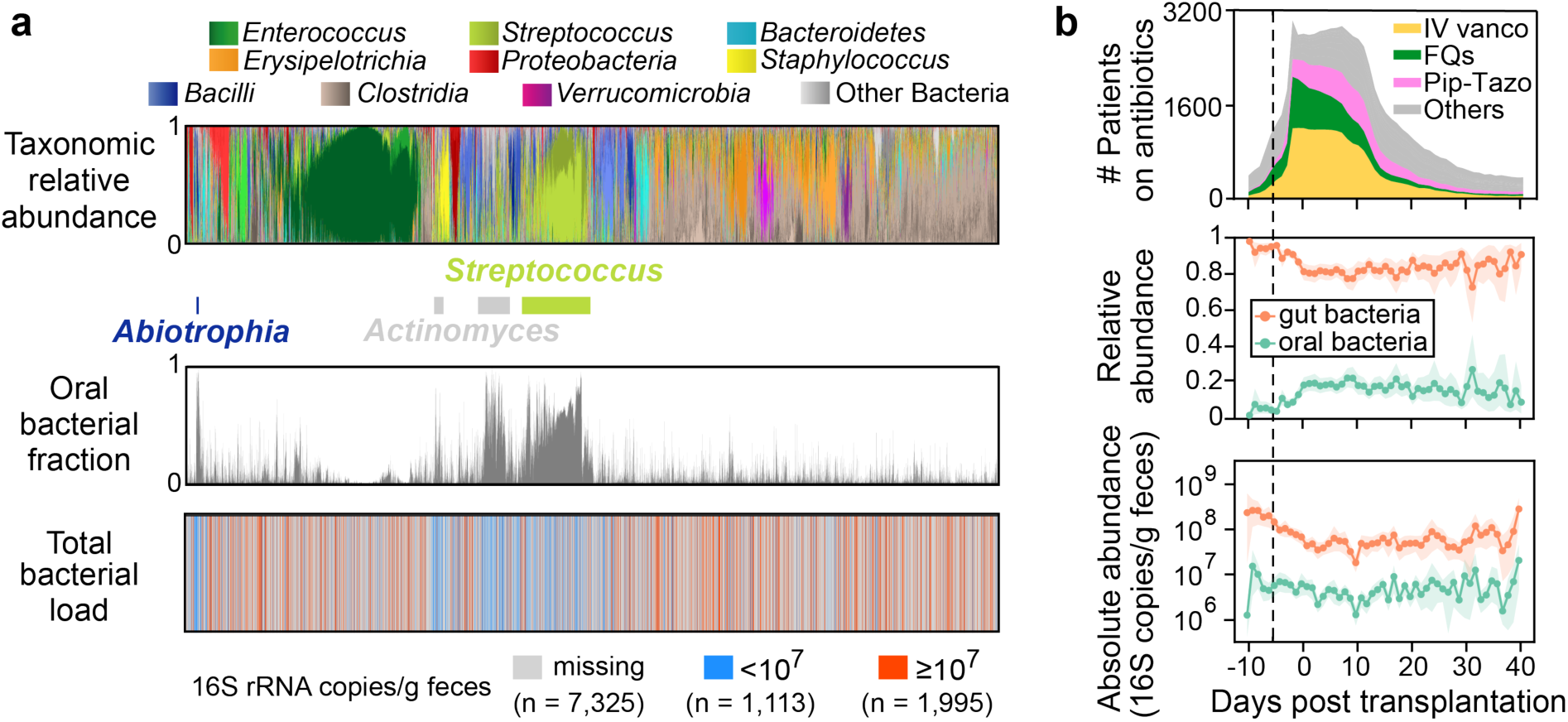
Relative enrichment of oral bacteria in fecal samples of allo-HCT patients coincides with onset of antibiotic prophylaxis and depletion of gut bacteria. **a**, Bacterial population composition in all 10,433 fecal samples from our MSKCC dataset. Each thin vertical bar represents a single sample, with taxonomic composition (top), estimated oral bacterial fraction (middle), and total bacterial load (bottom) aligned for all samples. **b**, Profiles of antibiotic exposure (top) aligned with the population dynamics of relative (middle) and absolute (bottom) abundance of oral (green curves) and gut (orange curves) bacteria in feces. Lines and dots represent the means, and shadings of the same color indicate the 95% confidence interval. The black dashed line marks the median day -6 when antibiotic prophylaxis was initiated. IV vanco: intravenous vancomycin; FQs: fluoroquinolones; Pip-Tazo: piperacillin-tazobactam.

As shown in Fig. 3b (top), oral bacteria became significantly enriched in the fecal samples before allo-HCT, which temporally coincided with the onset of antibiotic prophylaxis (Methods; n = 87, one-sided Wilcoxon signed-rank test, the same below; *P* = 1.2e-11, Cohen’s *d* = 1.09). Among the major antibiotics, piperacillin-tazobactam showed the most significant association with the enrichment (Table S5, Supplementary Note 4), which has been confirmed in a separate cohort of pediatric allo-HCT recipients (Fig. S2). Notably, the rise in the relative abundance of oral bacteria did not correlate with changes in their absolute abundance (Fig. 3b, middle and bottom, green curves), which remained stable without significant increase throughout the transplantation period (*P* = 0.364, Cohen’s *d* = 0.15). As a result, the relative enrichment was attributed to the significant reduction in the absolute abundance of gut bacteria (*P* = 9.9e-8, Cohen’s *d* = 0.65), as shown by the orange curve in the bottom panel of Fig. 3b.

Analogous to the analysis of our mouse experiment, we performed a linear regression analysis between the log_10_(oral bacterial fraction) and log_10_(total bacterial load). This analysis yielded a slope of -0.41 (Extended Data Fig. 6), which is higher than the theoretically expected value of -1 under the pure *Marker* hypothesis. However, by simulating the impact of interindividual variability on the regression slope (Methods), the theoretical value of the slope increased to -0.36 (Extended Data Fig. 7a), suggesting that the observed discrepancy may arise from the variability between patients. In contrast to the negative linear regression slope observed for oral bacteria, positive slopes were observed for several dominant gut bacterial genera, indicating that their relative abundance reflected their absolute abundance (Table S6).

We also analyzed metagenomic data available for a nested subset of 395 fecal samples^21^ (Methods). A peak-to-trough ratio analysis revealed a slow growth rate for the most abundant oral ASV (*Streptococcus* ASV_8), suggesting that the *Streptococcus* was not actively proliferating in the intestine (Table S7, Supplementary Note 5). Collectively, our human microbiome data analysis extends our findings beyond mice and validates the *Marker* hypothesis as the primary explanation for the enrichment of oral bacteria detected in patients’ feces.

### The *Marker* hypothesis is generalizable to IBD patients with depleted gut microbiota

The relative enrichment of oral bacteria in fecal samples has been recognized as a characteristic feature of IBD^22^. Intriguingly, another hallmark of IBD is a depleted microbial load in fecal samples^23,24^. If the *Marker* hypothesis holds for IBD patients, these two microbiome signatures should exhibit an inverse relationship. To test this hypothesis, we analyzed a previously published quantitative microbiome dataset obtained from 17 patients with Crohn’s disease and from 80 healthy controls^23^. These data included 16S amplicon sequencing and flow cytometric enumeration of microbial cells. As expected, these patients showed a higher oral bacterial fraction and lower total microbial load in their fecal samples compared to healthy controls (Fig. 4, marginal distributions). As a result, these two signatures were negatively correlated with a regression slope of -0.30, a value that matches the pure *Marker* hypothesis after considering interindividual variability (Extended Data Fig. 7b). We, therefore, conclude that the *Marker* hypothesis joins the two previously discovered IBD biomarkers and is generalizable to patients with Crohn’s disease.

**Figure 4:**
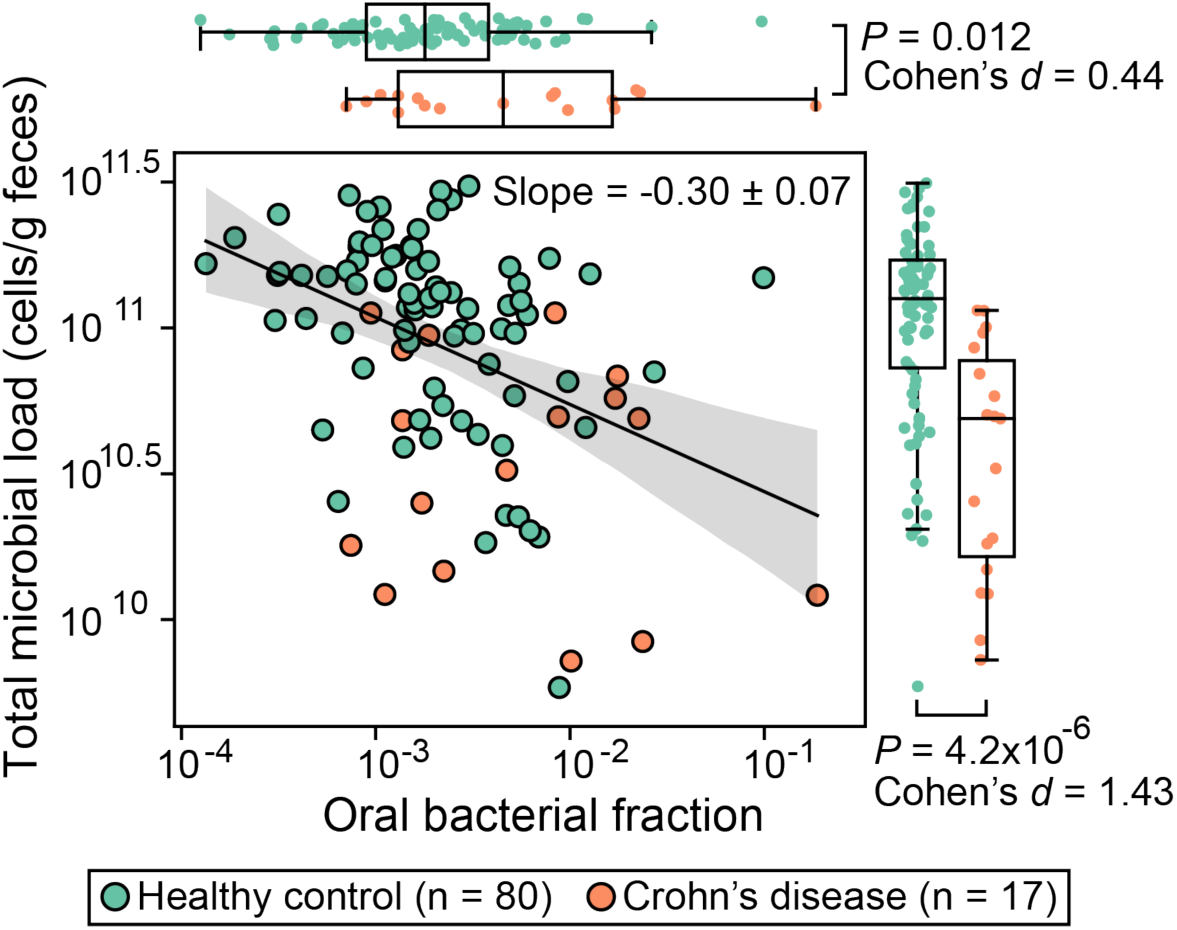
Inverse correlation between total fraction of oral bacteria and microbial load in fecal samples of patients with Crohn’s disease. Each circle represents a fecal sample. Samples with zero oral bacterial fraction in the original dataset were excluded from the plot and statistical analyses. The black line represents the best linear fit, and the shading of the same color indicates the 95% confidence interval. The plot also displays the regression slope and its standard error. *P* values for the difference in the marginal distributions between patient and control groups were calculated using a one-sided Mann-Whitney U test.

### Clinical implications of oral bacterial fraction as an indicator of total bacterial load

Loss of gut bacteria can alter host physiology and has been linked to host health^23–26^. Having established the *Marker* hypothesis, we sought to determine whether the oral bacterial fraction can replace the total bacterial load in assessing the impacts of gut bacterial depletion on patient outcomes. This substitution, if validated, has important clinical applications, since the associations between total bacterial load and patient outcomes could be effectively studied in the absence of quantitative microbiome data. To test the hypothesis, we used the MSKCC allo-HCT cohort which contains comprehensive patient outcome metadata. In this dataset, we showed that the oral bacterial fraction in feces can predict the depletion of total bacteria, with a mean cross-validation accuracy of 69% (Extended Data Fig. 8).

We considered four patient outcomes that can be influenced by gut bacterial depletion, including stool consistency, fungal overgrowth, bloodstream infection, and overall survival. If oral bacterial fraction could replace the total bacterial load, it should be associated with these patient outcomes. Stool consistency is strongly correlated with gut microbiome^27^. The loss of gut commensals reduces the total bacterial biomass and the production of short-chain fatty acids, which can increase water absorption in the colon^28^. As expected, we found a significant association between a high oral bacterial fraction and a low stool consistency (Extended Data Fig. 9a). Also, gut bacterial depletion can create an ecological niche for fungi to colonize and expand^29^. In line with our expectation, the oral bacterial fraction is significantly higher in fecal samples with a positive fungal culture than in those with a negative fungal culture (Extended Data Fig. 9b). Moreover, depletion of gut bacteria eliminates potentially invasive pathogens that may translocate to bloodstream and cause infections^30,31^. Indeed, our analysis revealed that a high oral bacterial fraction reduces the risk of total bacterial bloodstream infections (Methods; Hazard ratio = 0.29; 95% confidence interval, 0.11-0.75; *P* = 0.011). This result with a high oral bacterial fraction stands in contrast to intestinal domination states by *Enterococcus*, *Klebsiella*, or *Escherichia*, which strongly increase the risk of subsequent bloodstream infections caused by these organisms^30,31^. The striking contrast, in turn, supports our previous finding that the relative enrichment of oral bacteria in feces of allo-HCT recipients was not driven by their expansion in the gut (Fig. 3b). Finally, the oral bacterial fraction was negatively associated with all-cause and graft-versus-host disease (GVHD)-related mortality (Table S8, Supplementary Note 6), which is consistent with the notion that gut microbiome damage is associated with worse patient outcomes^32^.

## Discussion

The relative enrichment of oral bacteria detected in fecal samples has been associated with diseases^6^ and changes in host gene expression^13^ in various gastrointestinal disorders. To understand these associations, it is essential to distinguish changes in relative abundance from changes in absolute abundance of bacteria in feces^23,33^. In this study, we provide compelling evidence for the *Marker* hypothesis: a relative enrichment of oral bacteria signifies a depletion of commensals in the intestine rather than an expansion of the oral bacterial population with increasing absolute numbers. This conclusion is based on population averages. On an individual level, new oral ASVs may emerge in the gut following perturbations, leading to their increased absolute abundances (Extended Data Fig. 3). While we demonstrate that the total numbers of oral bacteria did not increase when gut bacteria have been depleted, it remains unclear what niche factors prevent their growth in the gut and whether these oral bacteria are even alive. Further research is needed to explore the growth and survival of oral bacteria in the gut niche^34^.

In our mouse experiments and those carried out in a previous study^17^, the administration of DSS-induced gut inflammation, leading to a drastic alteration of the gut environment and subsequent changes in gut microbiota composition. However, the chemically-induced gut inflamation did not reduce the total bacterial load in mouse feces. According to the *Marker* hypothesis, in such cases, an increase in the oral bacterial fraction should not occur due to the lack of gut bacterial depletion, which aligns precisely with our observations. Interestingly, the Crohn’s disease patients analyzed in Fig. 4 had lower levels of total bacterial load compared to the healthy controls, which correlated with a higher fraction of oral bacteria detected in their feces. This finding suggests that the mouse model of DSS-induced gut inflammation did not reproduce the gut bacterial depletion observed in these patients. Taken together, both the mouse and human data support the *Marker* hypothesis as the primary explanation for the observed gut microbiota changes in the context of gut inflammation.

Previous studies have shown that ectopic colonization of oral bacteria in mouse gut can induce strong immune responses and gut inflammation^3,5^. These findings in mice suggest a potential causal link between the presence of translocated oral bacteria in the gut and the development of diseases. By demonstrating the *Marker* hypothesis, our study suggests that these microbiome-disease associations could be mediated—at least in part—by the loss of gut commensals. In mice, the total microbial load in the intestine has exhibited positive correlations with fecal IgA concentration^24^, the proportions of mucosal RORγt^+^ cells^25^, and colonic lamina propria FoxP3^+^ regulatory T cells^24^. Still, the *Marker* hypothesis does not exclude the possibility that oral bacteria can drive disease. Even without an absolute increase in population size, oral commensals may acquire pathogenic potential in the human gut. Further research is needed to investigate how the pathogenicity of oral bacteria and the loss of gut commensals synergistically impact host responses and exacerbate pathology.

In summary, we have validated the *Marker* hypothesis by showing a robust negative correlation between oral bacterial fraction and total bacterial load under conditions where the gut microbiota is disrupted in mice, in patients undergoing allo-HCT, and in patients with inflammatory bowel disease. However, a limitation of our approach is that it depends on a marked depletion of gut bacteria caused by perturbations such as antibiotic treatments. For example, in our mouse experiment, antibiotics depleted the gut bacterial load by over 1,000-fold. When individuals exhibit minor or no variations in the size of their gut bacterial population^35^, the link between the fraction of oral bacteria detected in fecal samples and the total size of the gut bacteria population may be masked by interindividual variability. In such scenarios, the applicability of the *Marker* hypothesis and its derived results may also be limited.

Nonetheless, when subjects experience massive fluctuations in the total bacterial load, its negative correlation with oral bacterial fraction could provide a convenient method for estimating the changes in the absolute abundance of bacterial taxa from relative microbiome profiles alone. This can be achieved by calculating the ratio of their relative abundance to oral bacterial fraction in fecal samples (see Extended Data Fig. 10 for an example). This method could be applied to previously published microbiome compositional data to reevaluate previously identified bacterial taxonomic biomarkers and potentially discover new biomarkers based on estimated absolute abundances. Similarly, the oral bacterial fraction could serve as a biomarker for the outcome of fecal microbiota transplantation (FMT). A high fraction of oral bacteria in the feces of individuals being considered for FMT may signal a depleted gut microbiota in these candidates, thereby suggesting a higher chance of successful FMT engraftment. Supporting this notion, a recent study found that *Streptococcus salivarius*, a typical oral bacterial species, in the recipient’s microbiota facilitates donor strain colonization^36^.

## Acknowledgements

Chen.L. and A.D. were supported by National Institutes of Health (NIH) grant nos. U01 AI124275 (J.B.X.), R01 AI137269 (J.B.X.) and U54 CA209975 (J.B.X.). T.R. was funded by Deutsche Forschungsgemeinschaft (DFG, German Research Foundation) grant no. RO-5328/1-2 (T.R.), NIH grant nos. R01 AI093808 (T.M.H.), and R21 AI156157 (T.M.H.).

## Author contributions

Conceptualization, Chen.L., T.R., T.M.H., and J.B.X.; Mouse experiment, A.D., V.M., and Chloe.L.; Microbiome data processing, Chen.L., A.D., and H.L.; Microbiome data analysis, Chen.L., T.R. and T.F.; Writing – Original Draft, Chen.L., T.R., A.D., H.L.,T.F.; Writing – Review and Editing, T.M.H., J.B.X., J.U.P., B.Z., L.D., and M.R.M.v.d.B.; Supervision, J.B.X. and T.M.H.

## Data availability

A comprehensive list of the microbiome datasets used in this study, along with an indication of whether they contain quantitative data, is provided in Table S9. The raw 16S sequences of mouse fecal and oral samples collected in this study have been uploaded to Sequence Read Archive (SRA) with BioProject accession number PRJNA873058. All other microbiome datasets are publicly available. All processed data supporting the findings of this study are available within the article and its supplementary materials.

## Code availability

Customized Python codes for reproducing the figures and tables are available on Github (https://github.com/liaochen1988/Oral_bacterial_manuscript_revision)

## Methods

### Mouse experiment setup

Mice used in this study were C57BL/6J specific pathogen-free mice purchased from The Jackson Laboratories. Animals were 6-8 weeks old females. During the study, mice were single-housed in autoclaved cages with *ad libitum* access to autoclaved and acidified reverse osmosis water (pH, 2.5 to 2.8) and irradiated feed (LabDiet 5053, PMI, St Louis, MO). The animal holding room is maintained at 72 ± 2 °F (21.5 ± 1 °C), relative humidity between 30% and 70%, and a 12:12 hour light:dark photoperiod. Animal use is approved by MSKCC’s IACUC. The institution’s animal care and use program is AAALAC-accredited and operates in accordance with the recommendations provided in the *Guide*.

We performed two independent experiments. In the first one we tested the role of broad-spectrum antibiotics on the relative abundance of oral bacteria in the intestine and in the second one we repeated antibiotic cocktail treatment and included one more group that was exposed to dextran sulfate sodium (DSS) salt. Briefly, mice were treated with cocktail of ampicillin (0.5g/l), vancomycin (0.5g/l) and neomycin (1g/l) for one week in drinking water. To test the role of DSS, which is supposed to minimally affect bacterial load in the gut, we treated the mice with 3% DSS in the drinking water for a week. The water was changed once during the course of both treatments to preserve the activity of antibiotics and DSS. The third group of animals that served as a control drank regular water. Fecal pellets were collected at least immediately before and one week after the initiation of different treatments, and in some cases 3 days after the initiation of the treatment. Oral swabs were collected as per reference of Abusleme *et al.*^37^ at the time fecal sampling was done. Briefly, mice were hand-held while sterile swab was introduced into mouth and swiped for at least 30 seconds. After, the swab was put into 150 μl of TE buffer, and the tip was cut off so that the Eppendorf can be closed. Samples were put immediately to dry ice. One negative control swab was taken by pulling out the swab from the pouch and swirling through air for at least 30 seconds, after which it was put in TE and on dry ice. Fecal samples and oral swabs were kept at -80°C until further processing.

### DNA extraction and sequencing

DNA extraction and library preparation of the fecal samples collected during the first mouse experiment was done in the sequencing core facility at MSKCC. However, since the DSS treatment results in difficult amplification of the DNA, all fecal samples from the second experiment (control, antibiotic-treatment, and DSS-treatment samples) were subjected to DNA extraction and further processing in the Xavier lab with protocols adapted to ensure the proper DNA amplification and library preparation.

1) First experiment fecal samples processing. Fecal DNA was extracted and 16S rRNA gene was amplified using the previously described protocol^38^. QIAseq 1-step Amplicon Library Kit was used for generating libraries that were later quantified, normalized, and sequenced using MiSeq Reagent Kit V3.
2) Second experiment fecal samples processing. Bacterial DNA from fecal samples was extracted using the QIAamp® DNA Fast Stool Mini kit with introduction of a mechanic disruption step with bead-beating as per reference of Djukovic *et al.*^39^. The V4-V5 region of the 16S rRNA gene was amplified with Q5 High-Fidelity DNA Polymerase following the manufacturer’s recommendations. Briefly, for each sample, 25 μl of the PCR was prepared containing 5 μl of the Q5 buffer, 5 μl of the Q5 enhancer, 0.5 μl of the 10 mM dNTPs and 0.625 μl of the Q5 polymerase. 2.5 μl of the forward and 2.5 μl of the reverse primer at concentrations of 10 μM were added. Forward and reverse primers used for each sample contained unique sequences that would allow demultiplexing during sequence processing step. 1 ng of the DNA extracted from fecal samples collected from control and antibiotic-treated groups was added to reaction, while the DNA obtained from DSS-treated animals was diluted 1:20 and 5 ul was amplified. This was done since the carryover DSS prevented amplification of the DNA from some fecal samples, while in diluted samples the amplification worked as expected. Where necessary, the volume of the reaction was completed with water. Cycling conditions of the PCR were 96 °C for 10 minutes, and 35 cycles of 96 °C for 10 seconds, 51 °C for 30 seconds and 72 °C for 30 seconds. The final elongation step was performed at 72 °C for 5 minutes. Amplification was confirmed with agarose gel electrophoresis and PCR products were purified with AxyPrep Magnetic Beads, quantified with Quant-iT PicoGreen dsDNA kit, normalized and pooled. KAPA LTP Library Preparation Kit was used to generate sequencing libraries that were later quantified with Quant-iT PicoGreen dsDNA kit, normalized, and sequenced using MiSeq Reagent Kit V3.
3) Oral samples processing. Oral DNA was extracted by using modified DNeasy Blood and Tissue Kit protocol as described in Abusleme *et al.*^37^. After extraction V4-V5 region of the 16S rRNA gene was amplified with Q5 High-Fidelity DNA Polymerase by cycling 1 ng of the extracted DNA. Volumes and concentrations of PCR reagents were the same as described above. The following steps that include agarose gel electrophoresis, PCR product purification, normalization, pooling, library preparation, library quantification, and sequencing were performed as described in the previous section.

### Quantitative PCR (qPCR) for determining bacterial load

For assessing the bacterial load in fecal and oral samples, qPCR against standard curve was used to determine 16S rRNA copy number. For this purpose, the PowerUP qPCR Kit was used. Briefly, for each sample, 20 μl PCR triplicates were prepared with each containing 2 μl of the DNA used as template, 10 μl of mix provided by the manufacturer, and 1 μl of forward and reverse primers at the final concentration of 0.5 μM. We used the primer pair 27F/338R to amplify the V1-V2 region of the 16S rRNA gene (F-AGAGTTTGATCMTGGCTCAG; R-TGCTGCCTCCCGTAGGAGT). In order to complete the volume of the reaction, 6 μl of water was added. A PCR product of the 16S rRNA gene from *Enterococcus faecium* ATCC 700221 strain was used for obtaining a standard curve by amplifying its 16S rRNA gene and purifying the product. The copy number of the PCR product was determined based on its concentration and 16S rRNA sequence. A standard curve was obtained by using 10-fold dilutions.

Cycling conditions of the qPCR were 50 °C for 2 minutes, 95 °C for 2 minutes, and 40 cycles of 95 °C for 15 seconds, 56 °C for 15 seconds and 72 °C for 60 seconds. By extrapolating results by looking the ones obtained from standard curve samples, the number of 16S rRNA genes was determined for each sample. The final number of 16S rRNA genes per 1 g of fecal sample was calculated by multiplying the number of 16S rRNA molecules obtained by qPCR with DNA elution volume after DNA extraction and dividing this number with the weight of the fecal pellet from which DNA extraction was performed.

### 16S amplicon sequencing data processing

Samples collected from the first mouse experiment were sequenced once. Fecal and oral samples from the second mouse experiment were sequenced twice using the MiSeq sequencers at the Joao Xavier and Thomas Norman labs at MSKCC. Since the two sequencing runs yielded very similar read coverage and bacterial profiles, we combined the read counts from both runs for samples in the second mouse experiment. Sequence processing was done following DADA2 (Divisive Amplicon Denoising Algorithm) tutorial with an in-house script. Briefly, after demultiplexing, reads were trimmed to the first 180 bp or the first point with a quality score Q < 2, and removed if they contained ambiguous nucleotides (N) or if two or more errors were expected based on the quality of the trimmed reads. Paired reads were merged, and chimeras were removed. ASVs were identified using DADA2 and classified against the SILVA v138 database^40^.

The bacterial profiles of > 10,000 fecal samples from MSKCC allo-HCT recipients were previously analyzed by an in-house pipeline and compiled in a recent study^20^. Similarly, reads were trimmed to the first 180 bp or the first point with a quality score Q < 2, and removed if they contained ambiguous nucleotides (N) or if two or more errors were expected based on the quality of the trimmed reads. ASVs were identified using DADA2^41^ and classified against the SILVA v138 database^40^.

The demultiplexed and primer-trimmed HMP^1^ 16S sequences (V3-V5 region) were obtained from the Qitta repository^42^ and processed using QIIME (Quantitative Insights Into Microbial Ecology) 2^43^. DADA2 was applied to denoise the data and generate an ASV per sample count table, using the QIIME denoise-pyro plugin^43^. To remove low-quality tails, we used parameter --p-trunc-len 395 to generate ASVs that cover the entire high-quality V4-V5 region. Taxonomy classification of the ASV sequences was performed using the QIIME plugin “feature-classifier”^44^ and the SILVA v138 database^40^. The classification took three steps. We first extracted the V3-V5 region of the SILVA reference sequences using the extract-reads method. Then we created a classifier by using the fit-classifier-naïve-bayes method with extracted reads and the SILVA reference taxonomy. Finally, we ran the classifier on the ASV sequences using the classify-sklearn method to get their taxonomy.

All other microbiome datasets in this study were processed using QIIME 2^43^. Demultiplexed short reads were trimmed using the QIIME cutadapt plugin^45^ with parameters “-- p-error-rate 0.1” and “--p-overlap 3”. The trimmed reads were then denoised using the QIIME dada2 plugin with truncation lengths determined by per-base quality scores to generate ASV-level feature tables. Taxonomic classification was performed using the QIIME plugin “feature-classifier classify-sklearn”^46^ against the SILVA v138 database^40^.

For all microbiome datasets analyzed in this study, samples with sequencing coverage below 1,000 reads were excluded. Additionally, we excluded non-bacterial ASVs and ASVs with “Chloroplast” or “Mitochondria” in their taxonomy labels.

### Identification of oral ASVs in fecal samples

We leveraged paired oral and fecal samples to identify bacterial ASVs typically colonizing the oral cavity. An ASV is considered of oral origin if it satisfies all four of the following criteria simultaneously: (1) its relative abundance, averaged across all oral cavity samples, is above θ_*a*_; (2) its relative abundance, averaged across all fecal samples, does not exceed θ_*a*_; (3) its prevalence, across all oral cavity samples, is above θ_*p*_; (4) its prevalence, across all fecal samples, does not exceed θ_*p*_. We identified a total of 53 such oral ASVs in all mice using θ_*a*_ = 1e-3 and θ_*p*_ = 10%. Mouse-specific oral ASVs were selected by excluding any ASV from this set if it was undetectable in the pre-treatment oral samples. Exceptions were made for three mice (Control_2D, DSS_2A, DSS_2C) lacking pre-treatment oral samples, for which no ASVs were excluded. To identify oral ASVs in humans, we employed the HMP^1^ dataset and applied θ_*a*_ = 1e-4 and θ_*p*_ = 5%. The cutoff values are consistent with those used in a previous study^47^ for identifying oral species (not ASVs) from metagenomic data. For both mice and humans, the prevalence of an ASV was calculated as the proportion of samples containing the ASV at a relative abundance above 1e-3^48^. Unless otherwise specified, we calculated the absolute abundance of oral and gut bacteria by multiplying their relative abundances by the total bacterial loads, which can be quantified by either 16S qPCR or flow cytometry.

### Impact of interindividual variability on regression slope between oral bacterial fraction and total bacterial load

In the absence of interindividual variability, the derivative of the log-transformed total bacterial load with respect to the log-transformed oral bacterial fraction equals to -1 under the pure *Marker* hypothesis and remains positive under the pure *Expansion* hypothesis (see Supplementary Note 1). To understand the impact of interindividual variability on this derivative, we simulated the oral bacterial fraction (*f_oral_*) and total bacterial load (*F_total_*) in fecal samples from both a control and a case group. The control group samples are collected from individuals with low oral bacterial fraction. These individuals, for example, include healthy individuals or patients before a medicial treatment. The case group samples are collected after a perturbation that induces relative enrichment of oral bacteria, such as antibiotic treatment and certain intestinal disorders. It is important to note that the control and case group samples do not necessarily come from the same longitudinal study and may be collected from different populations in a cross-sectional study.

For each control group sample *i*, we assume that the log_10_-transformed values of the oral bacterial fraction (*f_oral,i,ctr_*) and total bacterial load (*F_total,ctr_*) follow separate Gaussian distributions:

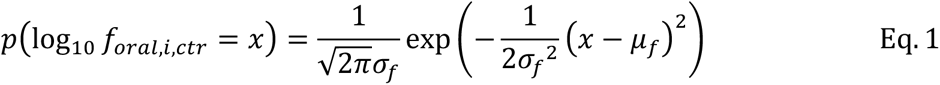

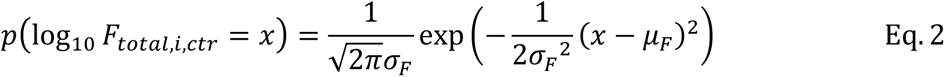

Here, μ_*f*_ and μ_*F*_ represent the means of the respective distributions, and σ_*f*_ and σ_*F*_ represent the standard deviations.

For each case group sample *j*, we first generate its baseline values of oral bacterial fraction (*f*_*oral,j,bl*_) and total bacterial load (*F*_*total,j,bl*_) from the same Gaussian distributions. These baseline values characterize the microbiome state prior to any perturation that enriches oral bacteria in this sample. The log_10_-transformed value of oral bacterial fraction after the perturbation (*f*_*oral,j,case*_) is drawn from a uniform distribution between log_10_ *f*_*oral,j,bl*_ and log_10_*f*_oral,max_. Here, *f*_*oral,max*_ represents the maximum possible oral bacterial fraction, which may vary based on the strength and type of the perturbation. After sampling *f*_*oral,j,case*_, the total bacterial load (*F*_*total,j,case*_) in this sample can be computed in two different ways. Under the pure *Marker* hypothesis (i.e., oral bacterial load remains constant), we have *F*_*total,j,case*_ = *f*_*oral,j,bl*_*F*_*total,j,bl*_/*f*_*oral,j,case*_. Under the pure *Expansion* hypothesis (i.e., gut bacterial load remains constant), we have *F*_*total,j,case*_= *F*_*total,j,bl*_(1 − *f*_*oral,j,bl*_)/(1 − *f*_*oral,j,case*_).

The simulation parameters (μ_*f*_, μ_*F*_, σ_*f*_, σ_*F*_) specific to the MSKCC allo-HCT cohort presented in Fig. 3 and the Crohn’s disease cohort presented in Fig. 4 were determined as follows. For the allo-HCT recipients, their fecal samples collected between day -20 and 40 relative to the day of transplantation were divided into two sets: 185 samples obtained prior to antibiotic prophylaxis constitute the control group, and 2,339 samples after the initiation of prophylaxis constitute the case group. Using the control group samples, we estimated the values of the following parameters: μ_*f*_ = -1.72, μ_*F*_ = 7.79, σ_*f*_ = 0.70, and σ_*F*_ = 0.99. On the other hand, the Crohn’s disease cohort consists of 80 healthy individuals in the control group and 17 patients with Crohn’s disease in the case group. We calculated the following parameter values from the control group samples: μ_*f*_ = -2.71, μ_*F*_= 11.01, σ_*f*_ = 0.48, and σ_*F*_ = 0.33.

### Cox proportional hazard model

We used a time-varying Cox proportional hazard model to evaluate the association between antibiotic exposure and intestinal domination by oral ASVs in allo-HCT recipients. A total of 291 patients with at least 10 samples between day -10 and 40 relative to transplantation were included. The event of interest was intestinal domination of any oral bacterial ASV that exceeds 30%. Binary indicators of antibiotic exposure were used as covariates (see Table S5 for included antibiotics). Patients without any oral bacterial domination at day 40 were censored. The model outputs the hazard ratio, which compares the likelihood of oral ASV domination occurring among patients with antibiotic exposure and patients without.

Using data from the same 291 patients, we performed the Cox proportional hazard regression analysis to assess the association between oral bacterial domination and total bloodstream infections (BSIs). Oral bacterial domination, as defined above, was analyzed as a time-varying predictor. The events of interest were BSIs caused by any bacterial species from *Citrobacter, Enterobacter, Enterococcus*, *Klebsiella*, *Escherichia*, *Pseudomonas*, *Stenotrophomonas*, and *Streptococcus*. Patients without any infection of interest at day 40 were censored. The hazard ratio indicates the relative risk of developing bacterial BSIs among patients with intestinal dominations of oral bacteria compared to those without dominations.

For both associations, we fit the Cox models using the CoxTimeVaryingFitter method from the Python package lifelines.

### Survival analysis of allo-HCT recipients

Following our previous approach^32,49^, we employed multivariable survival models with time-varying covariates to examine the associations between oral bacterial fraction in feces and all-cause or GVHD-related mortality among allo-HCT recipients. A total of 1,268 patients with fecal samples collected on or after day 0 relative to transplantation were included. The endpoint of interest was all-cause mortality or GVHD-related mortality until 2 years after allo-HCT, where patients who survived beyond 2 years were censored. Here, GVHD-related mortality was defined as death due to GVHD or after GVHD onset, without cancer relapse^50^ or other-cause mortality. Therefore, the GVHD-related mortality is the endpoint of interest with relapse and other-cause mortality as competition risks. Due to the different endpoint characteristics, we applied the Cox model and Fine-Gray model for all-cause mortality and GVHD-related mortality, respectively. Both models included oral bacterial fraction as the predictor, along with other covariates including log-transformed *Enterococcus* absolute abundance (a risk factor for both all-cause and GVHD-related mortality^51^), age, underlying disease, graft source, and conditioning intensity. Since qPCR data was available for fewer than a third of the samples, we used the *Enterococcus*-to-oral bacteria relative abundance ratio to estimate the absolute abundance of *Enterococcus* across all 10,433 samples. We have validated the estimation in Extended Data Fig. 10. The theoretical underpinning for this estimation is provided by the inverse correlation between oral bacterial fraction and total bacterial load: dividing by the oral bacterial fraction is equivalent to multiplying by the total bacterial load. The two survival analyses were performed using functions coxph and finegray from the R package survival, respectively.

### Shotgun metagenomic sequencing data processing

We adapted a recently published bioinformatic pipeline^52^ to assemble bacterial genomes from metagenomic data. The pipeline uses MEGAHIT^53^ to assemble contigs from short reads. It subsequently employs Metabat2^54^ and CONCOCT^55^ to bin these contigs into Metagenome-Assembled Genomes (MAGs). In the last step, it uses DAS Tool^56^ to generate an optimized, non-redundant set of MAGs. High-quality *Streptococcus* MAGs (≥ 75% complete, ≤ 175 fragments/Mbp sequence, and ≤ 2% contamination) classified by Kraken2^57^ were further analyzed by iRep^58^. The iRep value of a MAG represents the average number of replication events over different subpopulations of the MAGs weighted by their relative abundances.

### Statistical testing

Detailed descriptions of statistical analyses, including sample size, hypothesis test name, *P* value, and effect size, are provided. All *P* values were corrected for multiple testing using false discovery rate (FDR) adjustment (the Benjamini-Hochberg procedure). Statistical significance was defined as (adjusted) *P* value < 0.05. The pairwise Adonis test was performed using the R package pairwiseAdonis. The Mann-Whitney U test, Wilcoxon signed-rank test, and linear regression analysis were performed using the corresponding functions in the Python Scipy package. Specifically, we used a one-sided Wilcoxon signed-rank test to assess the differences in the oral bacterial fraction, oral bacterial load, and gut bacterial load before and after the administration of antibiotic prophylaxis among MSKCC allo-HCT recipients. We defined the pre-treatment period as spanning from 20 days before transplantation until the onset of antibiotic prophylaxis. The post-treatment period extended from the initiation of antibiotic prophylaxis to the day of neutrophil engraftment. We computed their average values across all samples collected during each of the pre-treatment and post-treatment periods.

### Plots

For all main and supplementary figures displaying a boxplot, the central line represents the median. The box limits correspond to the first and third quartiles (25th and 75th percentiles). The whiskers extend to the smallest and largest values or at most to 1.5 times the interquartile range, whichever is smaller. All plots were generated using Python Matplotlib library and the Seaborn package.

---

**Extended Data Figure 1:**
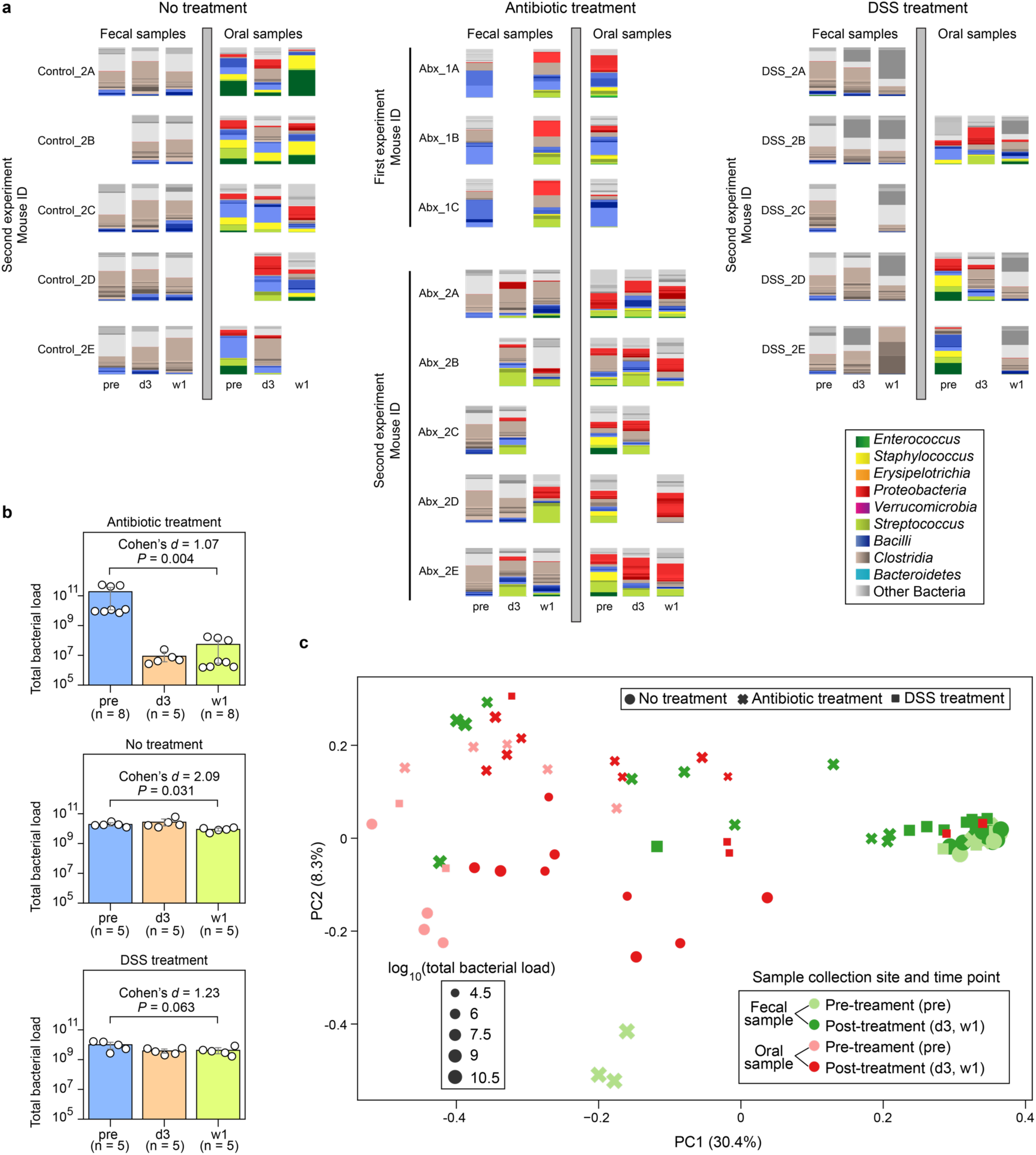
Paired fecal and oral microbiome samples in the mouse experiment. **a,** Bacterial ASV profiles displayed using stacked bar plots. Samples were collected at three time points: pre (before treatment), d3 (3 days after treatment initiation), and w1 (one week after treatment initiation). Missing samples in the second experiment were due to low sequence coverage (samples with less than 1,000 reads were excluded). **b**, Total bacterial load in fecal samples (circles). Bar heights represent the means, with error bars indicating the 95% confidence interval. *P* values were calculated using a one-sided Wilcoxon signed-rank test. **c**, Principal coordinate analysis plot of these samples. PC1 and PC2 represent principal component 1 and 2, respectively. The unit of total bacterial load is 16S copies per gram of feces for fecal samples and 16S copies per swab for oral samples. DSS: Dextran Sulfate Sodium.

**Extended Data Figure 2:**
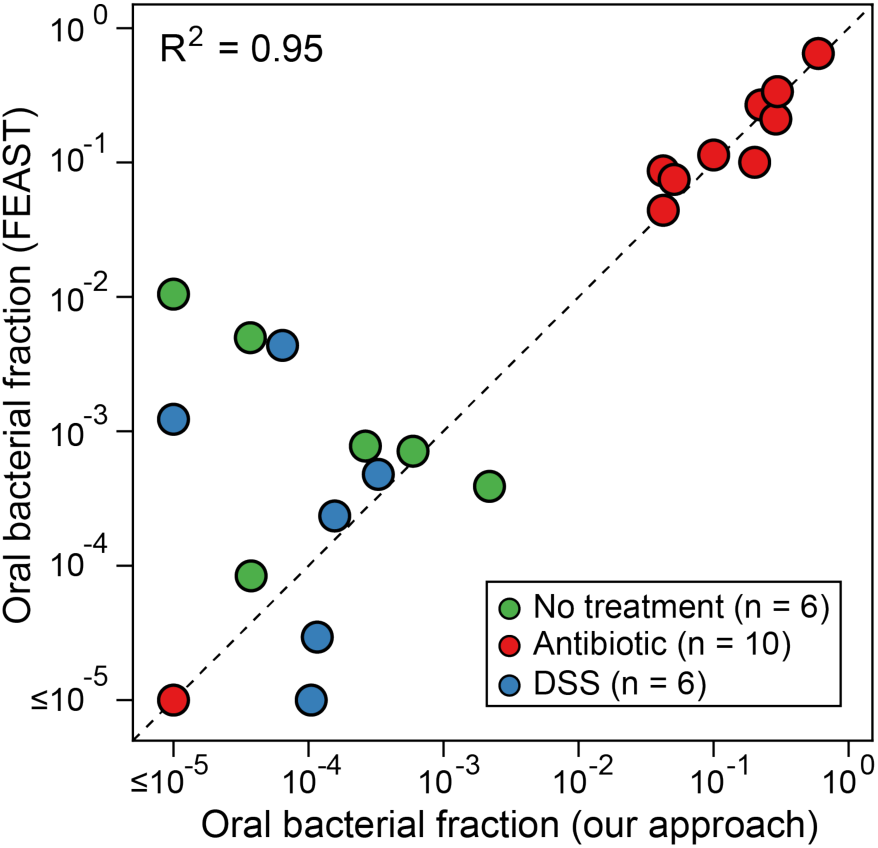
Quantitative agreement between our method and FEAST^18^ for estimating the oral bacterial fraction in post-treatment fecal samples. DSS: Dextran Sulfate Sodium.

**Extended Data Figure 3:**
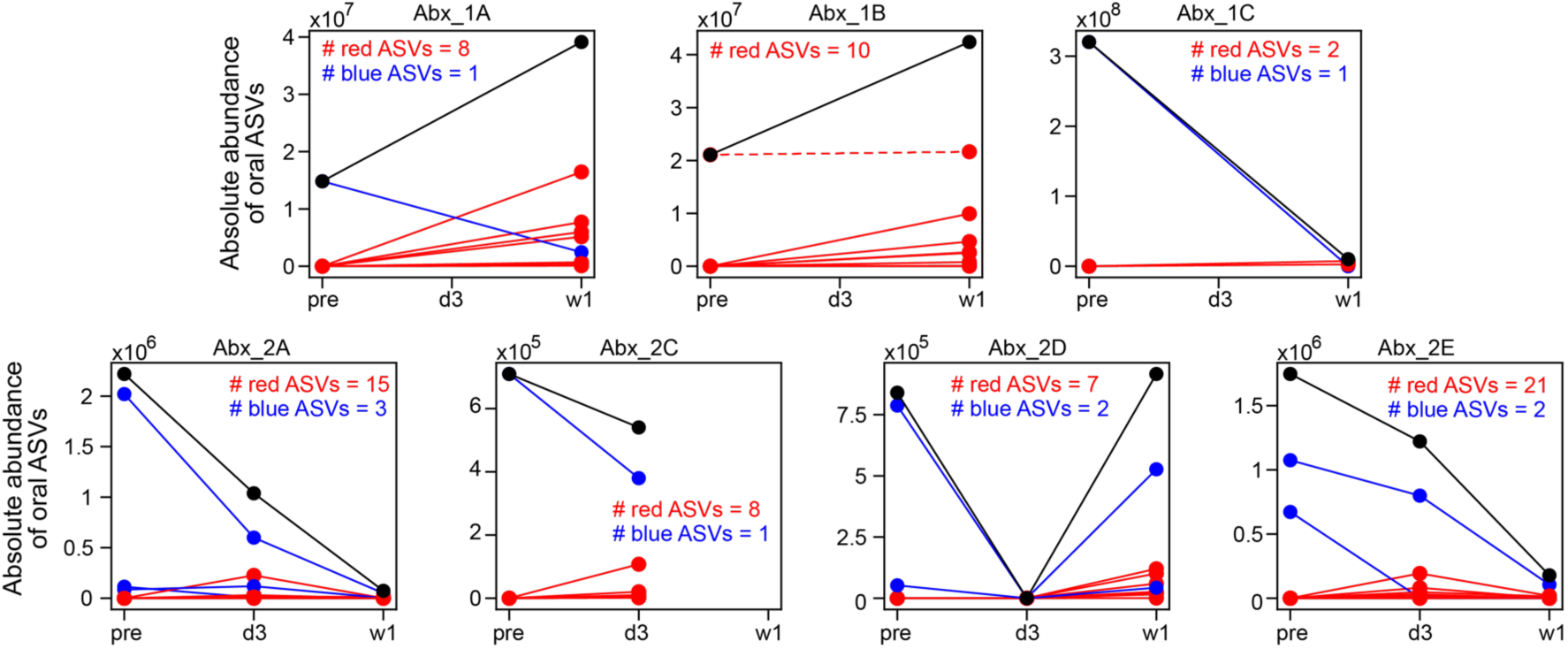
Emergence of new oral ASVs in mouse feces after antibiotic treatment. Each panel plots data from one mouse. Mouse Abx_2B was excluded due to the absence of its pre-treatment fecal sample. In each panel, individual oral ASVs are shown by lines. ASVs that increased in absolute abundance after treatment (i.e., post-treatment average > pre-treatment) are shown in red, while those that decreased or remained unchanged (post-treatment average σ; pre-treatment) are shown in blue. The black line in each panel represents the total absolute abundance, which is the sum of all red and blue lines. Except for one oral ASV in mouse Abx_1B (shown as a red dashed line), all ASVs displaying increased absolute abundance (solid red lines) were not present in the pre-treatment fecal samples; they represent new ASVs emerging from the oral cavity. Samples were collected at three time points: pre (before treatment), d3 (3 days after treatment initiation), and w1 (one week after treatment initiation). The unit for absolute abundance of oral ASVs in mouse feces is 16S copies per gram of feces.

**Extended Data Figure 4:**
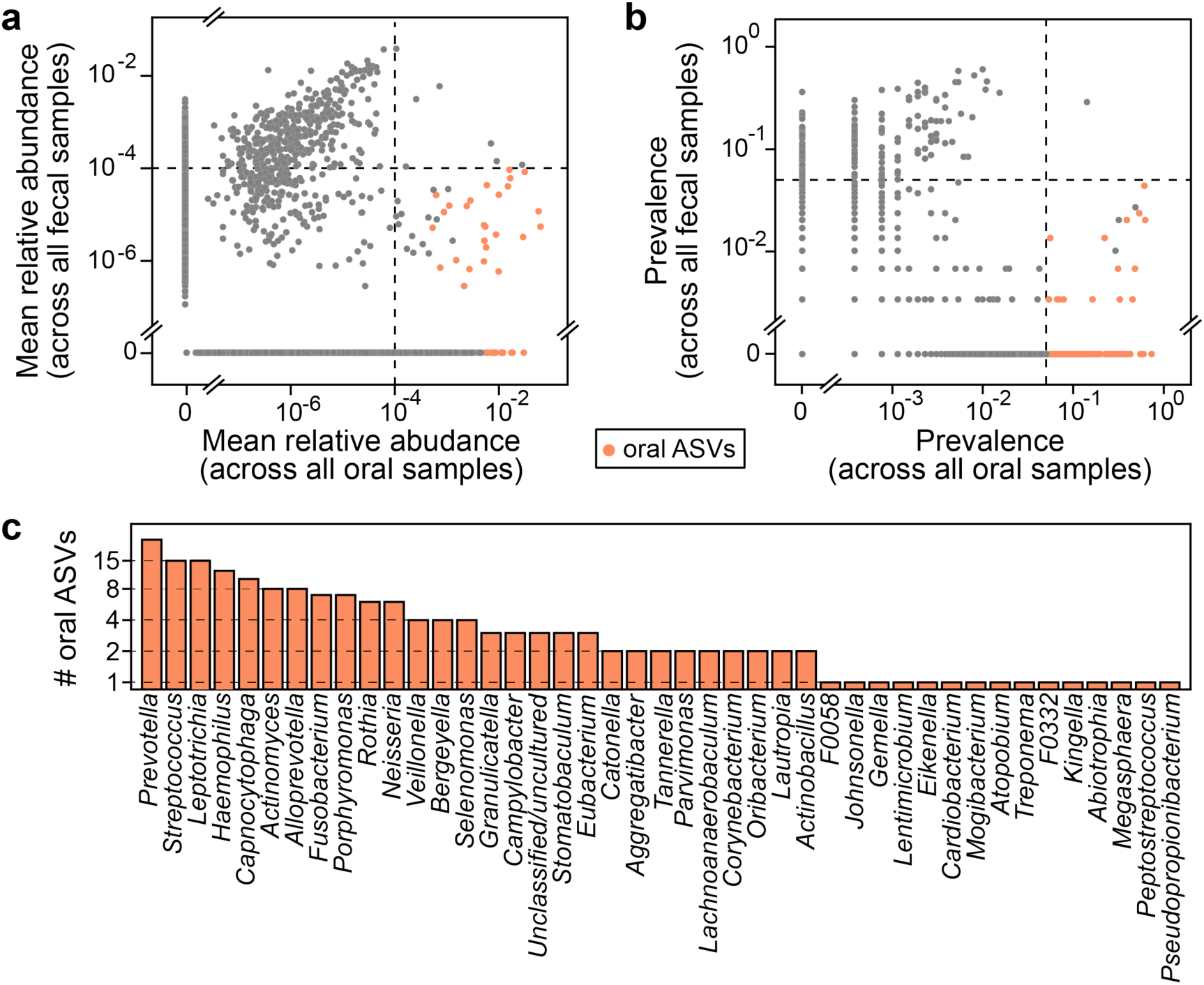
Oral bacterial ASVs identified from healthy human individuals involved in the HMP dataset. **a**,**b,** Mean relative abundance (a) and prevalence (b) of ASVs across 2,641 oral cavity samples (*x*-axis) and 291 paired fecal samples (*y*-axis). Black dashed lines represent the cutoffs used to identify oral ASVs (see Methods for details). Each dot represents an ASV. There is a total of 23,411 ASVs, where 178 ASVs identified as of oral origin are highlighted in orange. **c**, Distribution of the 178 oral ASVs at the genus level.

**Extended Data Figure 5:**
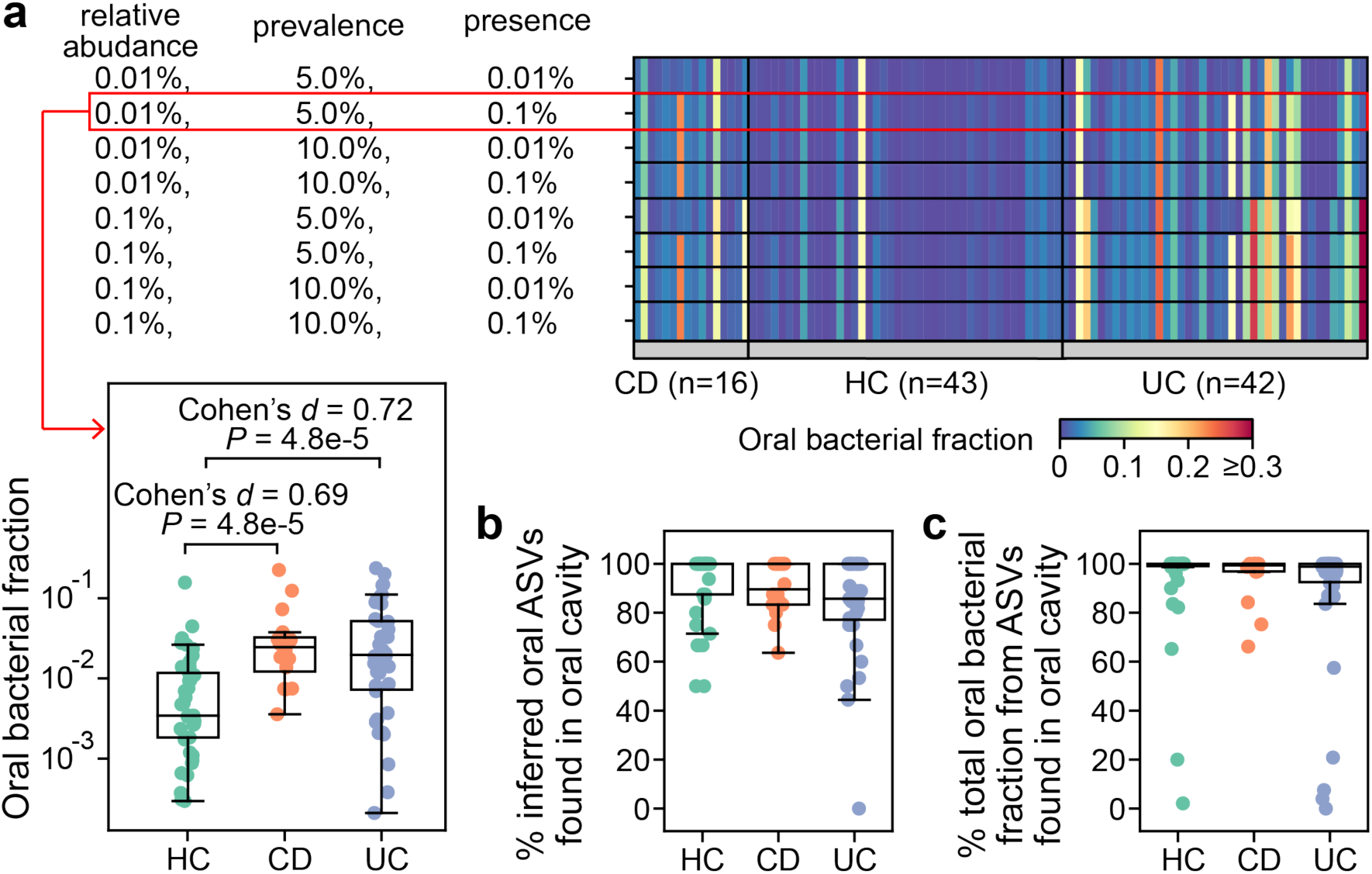
Validation of oral bacterial ASVs identified from healthy individuals in patients with inflammatory bowel disease. This cohort consists of 16 patients with Crohn’s disease (CD), 42 patients with ulcerative colitis (UC), and 43 healthy controls (HC). Both microbiome data and clinical metadata were obtained from a literature study^19^. Notably, none of the participants had taken antibiotics within the last three months prior to the study entry. **a,** Impact of cutoff parameters on the estimated oral bacterial fraction in feces of these participants. We systematically varied cutoffs for the mean relative abundance (0.01%, 0.1%), prevalence (5%, 10%), and the definition of ASV presence (0.01%, 0.1%) in the computation of prevalence to establish the reference set of oral ASVs. For each reference set, oral ASVs in their feces were inferred through exact sequence matching. The default parameter combination used throughout the study is outlined in the red box. *P* values were calculated using a one-sided Mann-Whitney U test. **b**, Percentage of inferred oral ASVs in feces that are also found in paired saliva samples. **c**, Percentage of the total relative abundance of inferred oral ASVs in feces contributed by those found in paired saliva samples. Two HC samples with a zero oral bacterial fraction are not shown in all box plots.

**Extended Data Figure 6:**
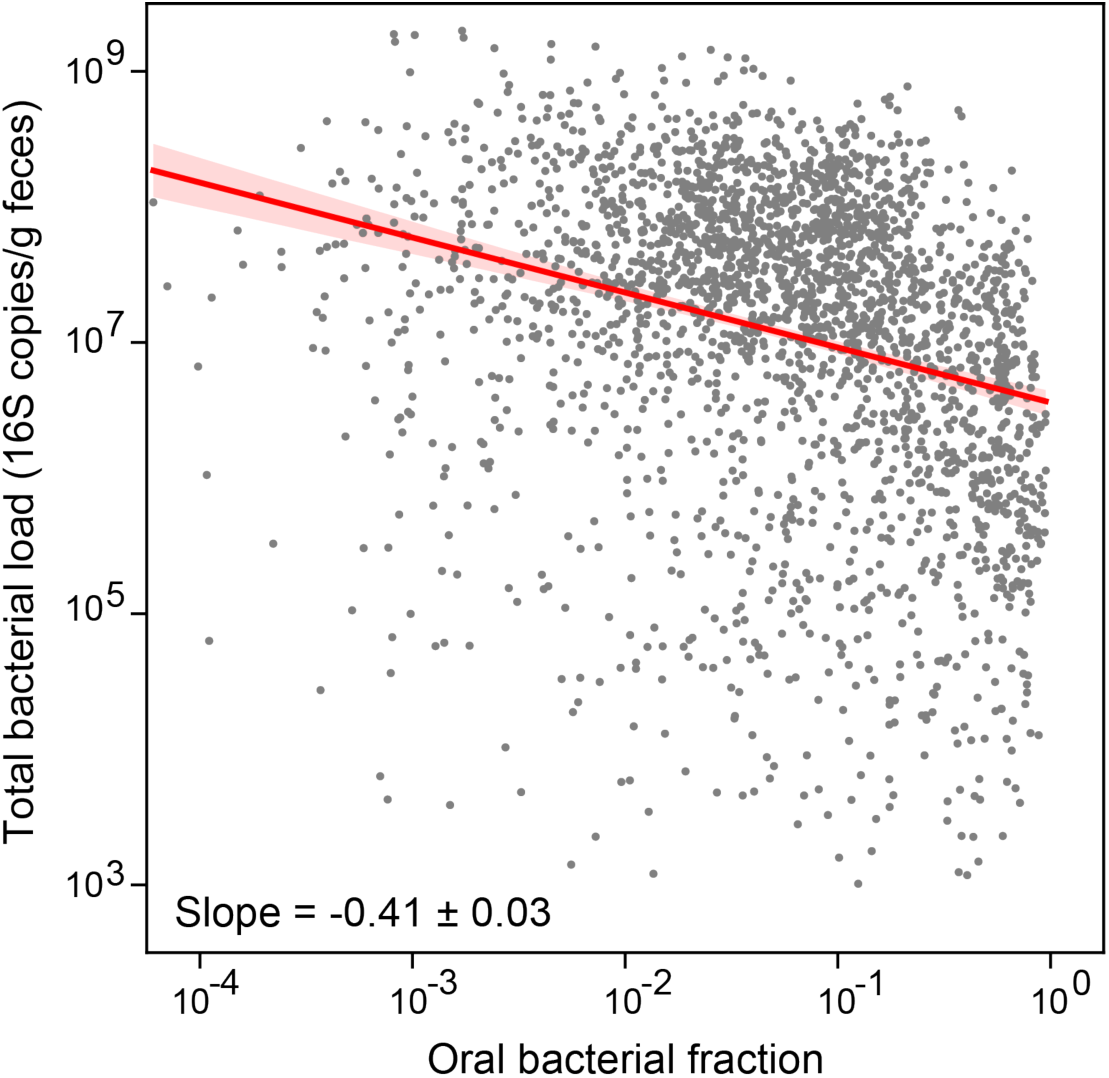
Inverse correlation between oral bacterial fraction and total bacterial load in fecal samples from MSKCC allo-HCT recipients. Each dot in the plot represents a fecal sample. Samples with zero oral bacterial fraction, or with total bacterial load less than 1,000 16S copies per gram of feces, or collected 20 days before or 40 days after transplantation were excluded from the plot and the linear regression analysis. The number of remaining samples is 2,524. The red line represents the best linear fit, and the shading of the same color indicates the 95% confidence interval. The plot also displays the estimated regression slope and its standard error.

**Extended Data Figure 7:**
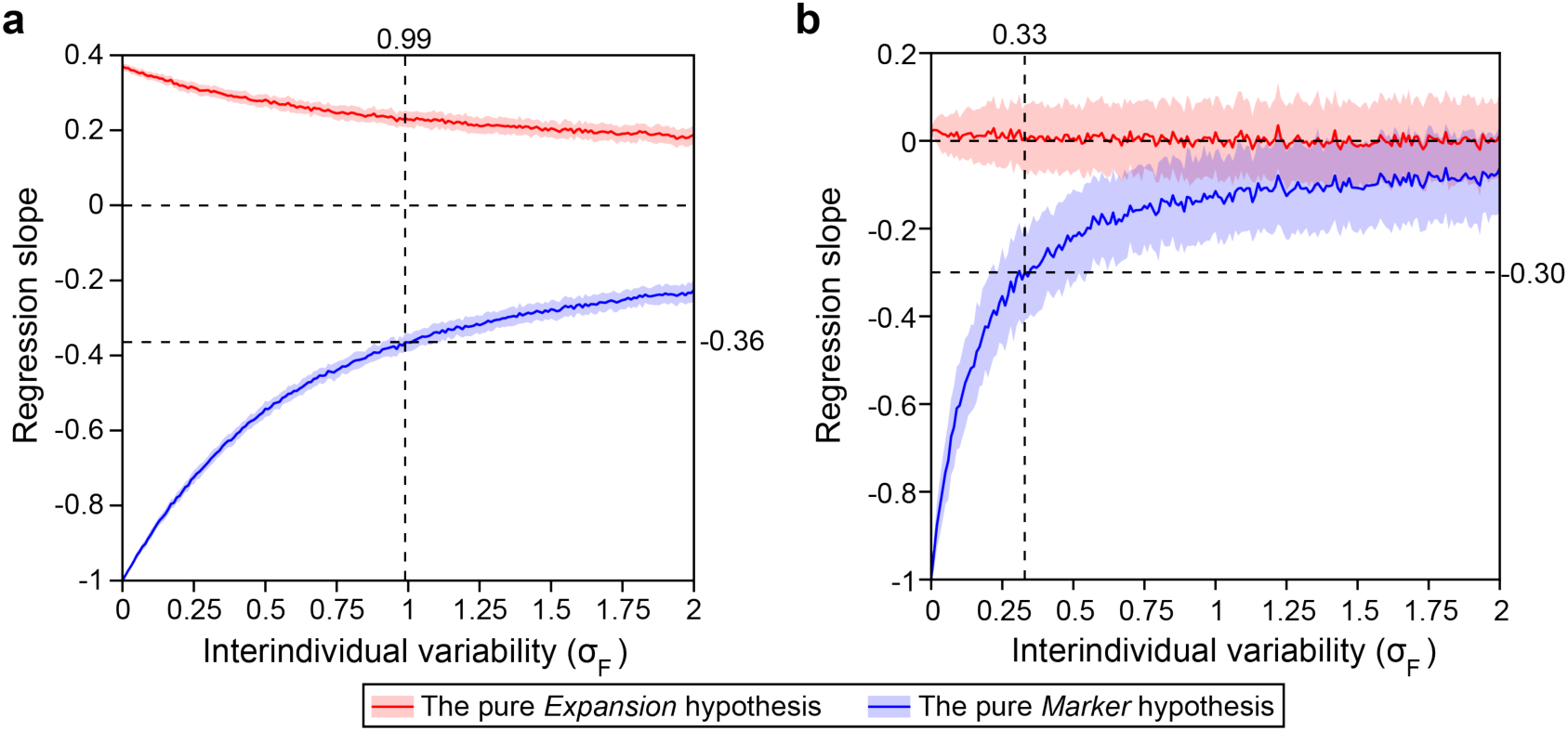
Impact of interindividual variability on the regression slope between oral bacterial fraction and total bacterial load. We generated synthetic datasets using parameters estimated from the MSKCC allo-HCT cohort analyzed in Fig. 3 (**a**) and the Crohn’s disease cohort analyzed in Fig. 4 (**b**). In both panels, we systematically varied σ_*F*_ (Eq. 2) from 0 to 2 while co-varying σ_*f*_ (Eq. 1) to maintain a constant ratio of σ_*f*_/σ_*F*_. For each synthetic dataset, the regression slope was determined through linear regression between oral bacterial fraction and total bacterial load in the log-log space. The red and blue lines represent the mean slopes over 100 simulation runs, with the shaded regions of the same color indicating the standard deviations. Vertical dashed lines in the panel (a) and (b) mark the σ_*F*_ values estimated from the pre-antibiotic-prophylaxis samples in the allo-HCT cohort and from the healthy individuals in the Crohn’s disease cohort, respectively. At these σ_*F*_ values, the pure *Marker* hypothesis predicted that the regression slopes are -0.36 and -0.30. For detailed information on our simulation approach, please refer to the Methods section.

**Extended Data Figure 8:**
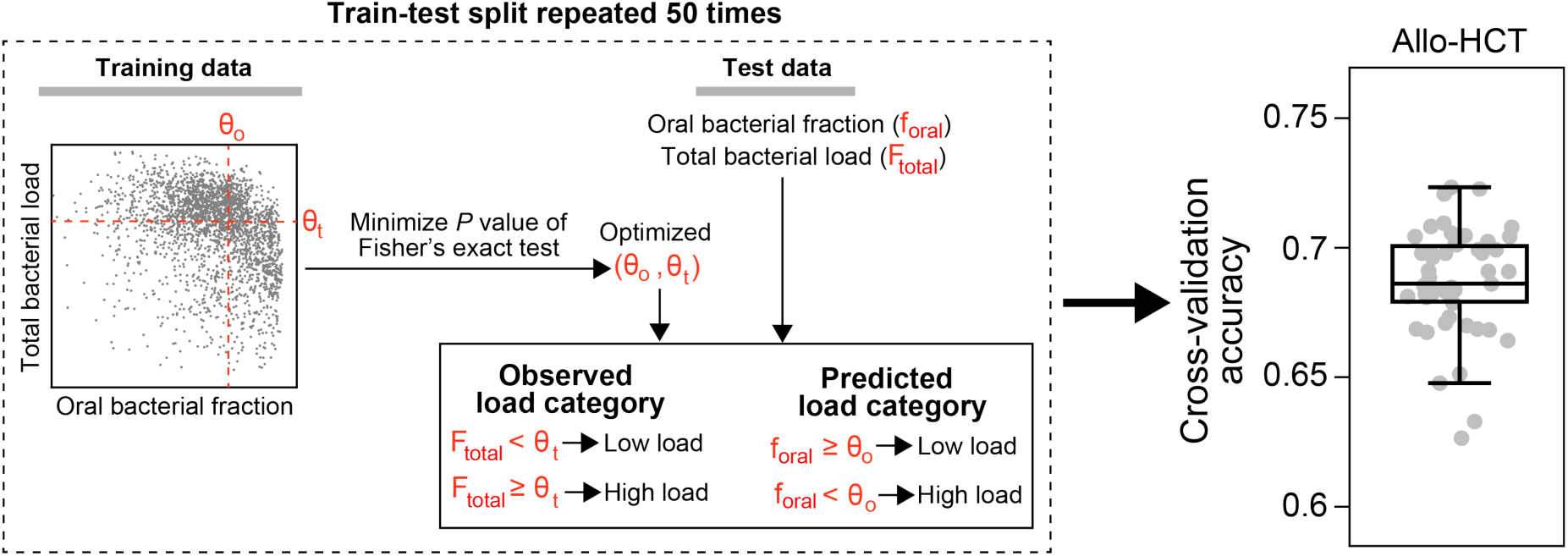
Cross-validation accuracy for classifying total bacterial load in fecal samples from MSKCC allo-HCT recipients. A schematic illustration of our classification model is enclosed in the dashed box. Our model uses two threshold parameters, θ_*o*_ and θ_*t*_, to convert the oral bacterial fraction and total bacterial load into binary categories, respectively. Given training data, we optimized the two cutoff parameters by minimizing the *P* value of Fisher’s exact test of independence, subject to the constraints θ_*o*_ ∈ [10^-4^, 1] and θ_*t*_ ∈ [10^3^, 10^10^]. The optimized θ_*o*_ was subsequently applied to predict high or low bacterial loads in the test set by comparing the oral bacterial fractions to θ_*o*_. Simultaneously, we binarized the observed bacterial loads by comparing them to θ_*t*_. Accuracy was assessed by comparing the predicted bacterial load categories to the observed bacterial load categories. In the boxplot, each dot corresponds to a single 5-fold cross-validation split and the random train-test split was repeated 50 times.

**Extended Data Figure 9:**
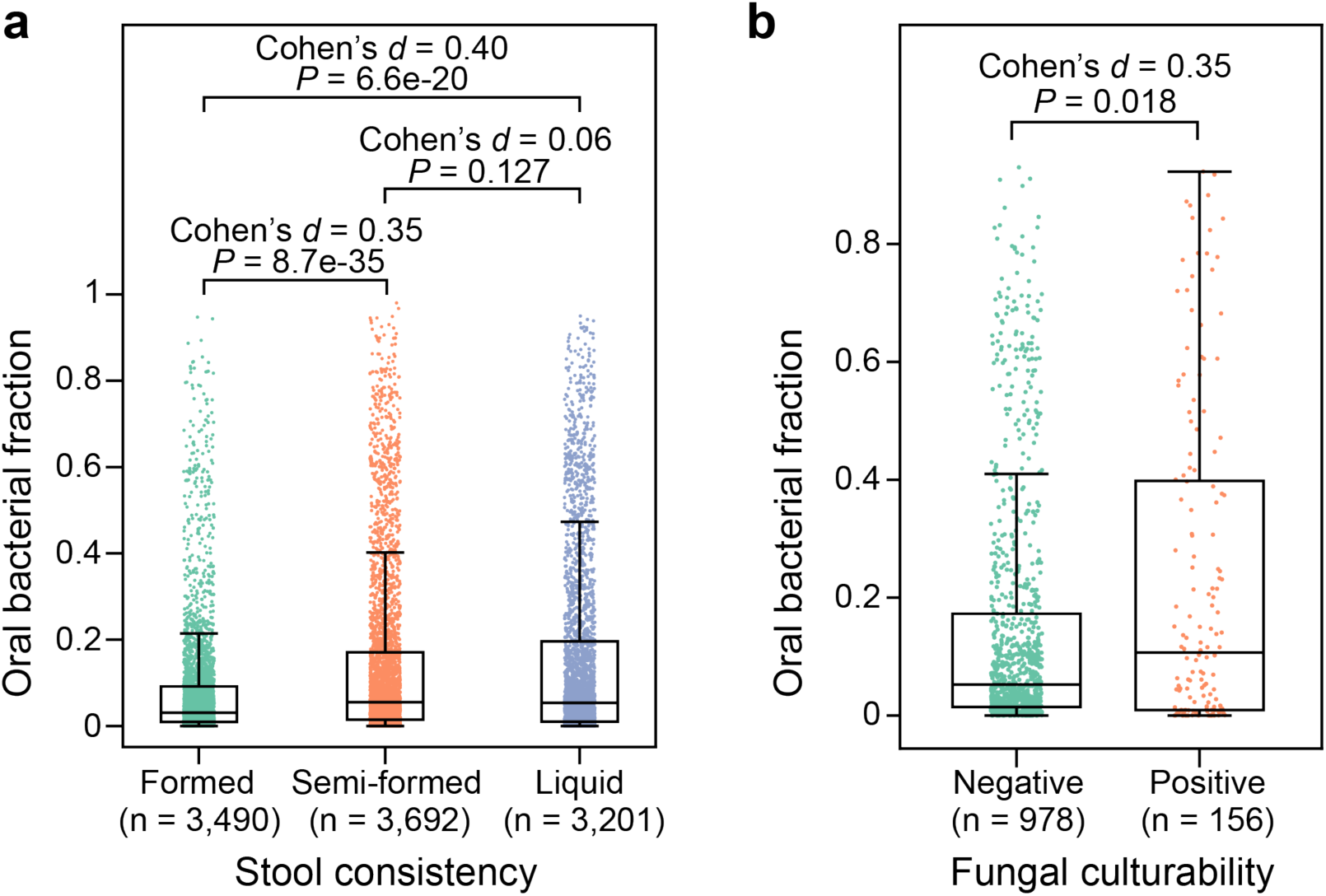
Distribution of oral bacterial fraction in fecal samples from MSKCC allo-HCT recipients, categorized by stool consistency (a) and fungal cultivability (b). Each dot represents a fecal sample. FDR-corrected *P* values were calculated using a two-sided Mann-Whitney U test.

**Extended Data Figure 10:**
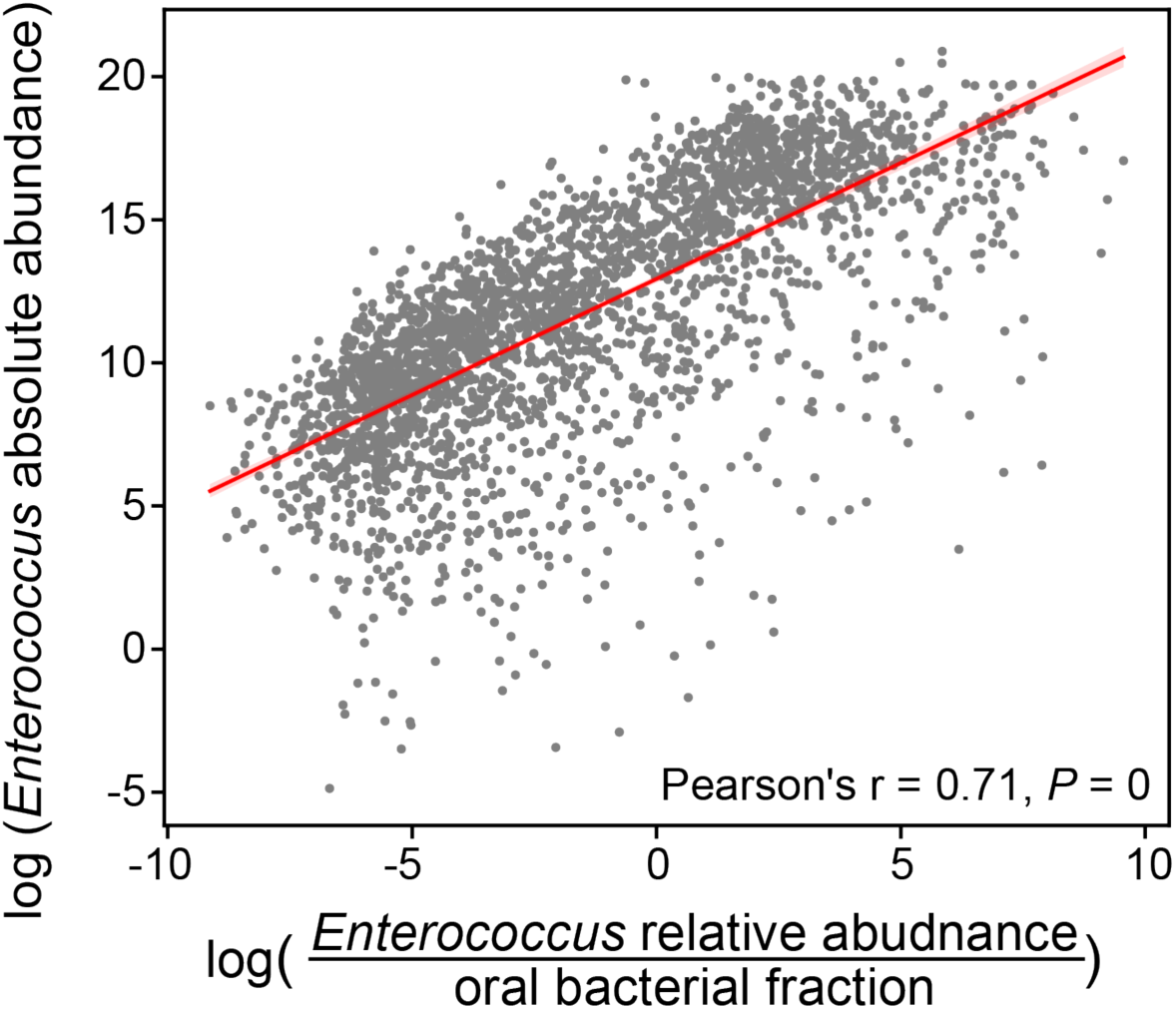
Estimating *Enterococcus* absolute abundance through the *Enterococcus*-to-oral bacteria relative abundance ratio. Each data point represents a fecal sample from MSKCC allo-HCT recipients. We excluded samples with zero relative abundance of *Enterococcus* or zero oral bacterial fraction from both the plot and the Pearson’s correlation analysis, resulting in a subset of 2,765 samples. The red line represents the best linear fit, with the shading in the same color indicating the 95% confidence interval. The absolute abundance of *Enterococcus* on the *y*-axis was calculated by multiplying its relative abundance by the total bacterial load measured by 16S qPCR, and thus has a unit of 16S copies per gram of feces.

