## Supplementary Information for "A High Fraction of Oral Bacteria in the Feces Indicates Gut Microbiota Depletion with Implications for Human Health"

**Liao *et al.***

**This PDF file includes:**

Figures S1, S2

Tables S2, S5, S8

Note: Tables S1, S3, S4, S6, S7, S9 are provided separately as excel files.

Supplementary Notes

Supplementary References

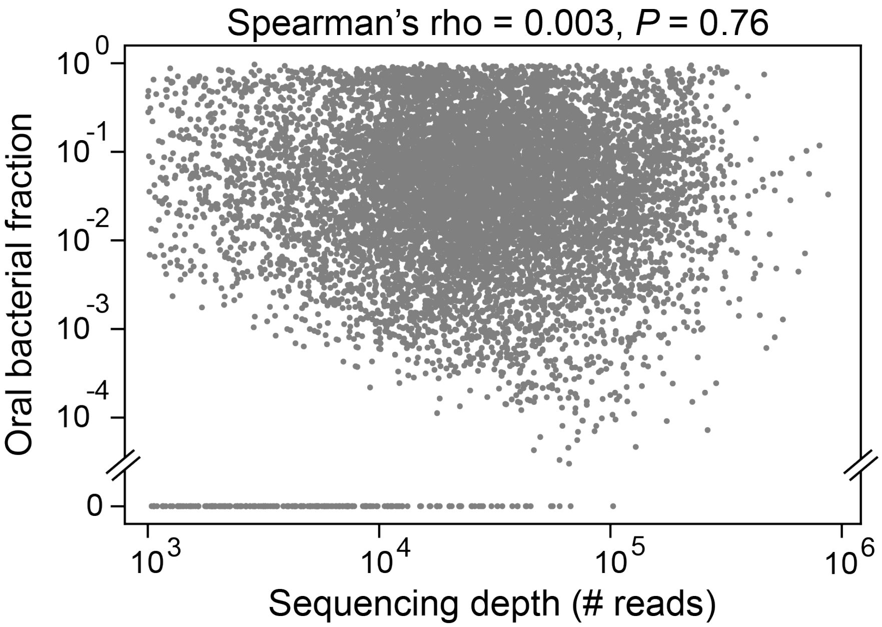


**Figure S1: No association between oral bacterial fraction and sequencing depth across fecal samples of MSKCC allo-HCT recipients.** Each dot represents a fecal sample (n = 10,433).

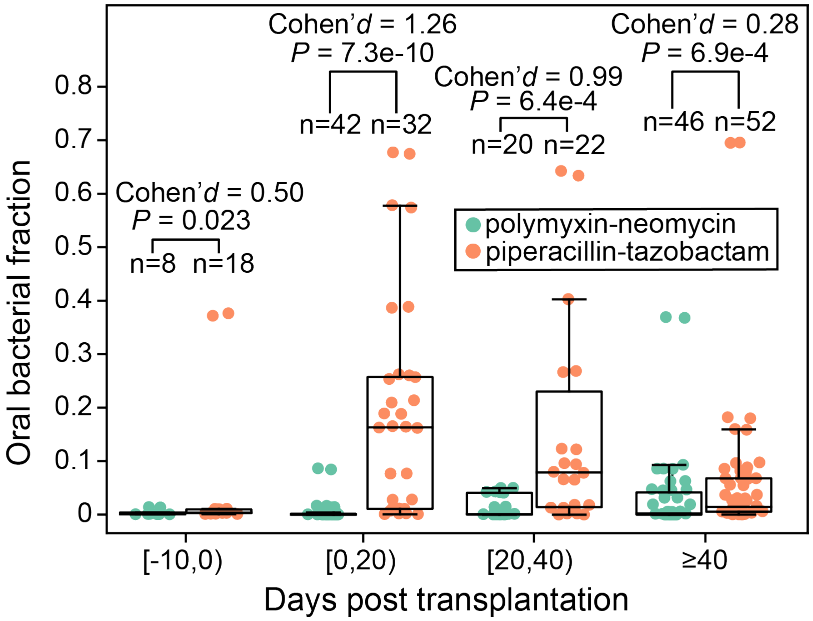


**Figure S2: Validation of piperacillin-tazobactam's effect on enriching oral bacteria in feces using an independent (non-MSKCC) pediatric allo-HCT cohort[1]**. 19 children (1-17 year old, 10.1 year old on average) were treated with either oral polymyxin-neomycin or oral piperacillin-tazobactam in the Leiden University Medical Center, Netherlands. Both medications were administered 10 days before transplantation until engraftment or 21 days after transplantation, whichever occurred later. Samples were grouped into four transplantation stages. FDR-corrected *P* values were calculated using a one-sided Mann-Whitney U test.

**Table S1: Oral bacterial ASV, taxonomy, and total abundance in mouse feces. a**, Sequence and taxonomy of all 53 oral bacterial ASVs found in mouse feces, with specific oral ASVs for each mouse indicated in the table. **b**, Total relative and absolute abundances of oral and gut bacteria in fecal samples. We determined the oral or gut bacterial load (absolute abundance) by multiplying the total bacterial load by the respective oral or gut bacterial fraction (relative abundance). Bacterial load units for fecal and oral samples are 16S copies per gram of feces and per swab, respectively. Samples with fewer than 1,000 reads were excluded from analysis and are labelled in the "Exclusion" column. DSS: Dextran Sulfate Sodium.

See separate Excel table

**Table S2: Pairwise Adonis test of compositional similarity between pre- and post-treatment fecal and oral microbiome samples.** Adonis test is a permutation-based multivariate analysis that assesses the differences in community composition between groups of microbiome samples. In our analysis, microbiome samples were divided into four groups: pre-treatment oral samples (group label: PreOral), pre-treatment fecal samples (group label: PreFecal), post-treatment oral samples (group label: PostOral), and post-treatment fecal samples (group label: PostFecal). The table below displays the $R^2$ values and FDR-corrected $P$ values for all six pairs of comparisons and for each mouse group. Bray-Curtis distance metric was used to measure the similarity between microbiome samples. Notably, post-antibiotic-treatment fecal samples exhibited greater similarity to pre-treatment oral samples ($R^2 = 0.15$, $P = 0.055$) than to pre-treatment fecal samples ($R^2 = 0.20$, $P = 0.032$).

| **Mouse group** | **Sample group 1** | **Sample group 2** | **$R^2$** | ***P* value** |
|---|---|---|---|---|
| No treatment | PreOral (n = 4) | PostOral (n = 9) | 0.21 | 0.058 |
| | PostOral (n = 9) | PostFecal (n = 10) | 0.76 | 0.001 |
| | PreFecal (n = 4) | PostOral (n = 9) | 0.71 | 0.003 |
| | PreOral (n = 4) | PostFecal (n = 10) | 0.92 | 0.003 |
| | PreOral (n = 4) | PreFecal (n = 4) | 0.95 | 0.044 |
| | PreFecal (n = 4) | PostFecal (n = 10) | 0.09 | 0.305 |
| Antibiotic treatment | PreOral (n = 8) | PostOral (n = 8) | 0.14 | 0.066 |
| | PostOral (n = 8) | PostFecal (n = 12) | 0.12 | 0.066 |
| | PreFecal (n = 7) | PostOral (n = 8) | 0.42 | 0.002 |
| | PreOral (n = 8) | PostFecal (n = 12) | 0.15 | 0.055 |
| | PreOral (n = 8) | PreFecal (n = 7) | 0.33 | 0.004 |
| | PreFecal (n = 7) | PostFecal (n = 12) | 0.20 | 0.032 |
| Dextran sulfate sodium treatment | PreOral (n = 3) | PostOral (n = 5) | 0.41 | 0.086 |
| | PostOral (n = 5) | PostFecal (n = 9) | 0.10 | 0.273 |
| | PreFecal (n = 5) | PostOral (n = 5) | 0.26 | 0.055 |
| | PreOral (n = 3) | PostFecal (n = 9) | 0.56 | 0.029 |
| | PreOral (n = 3) | PreFecal (n = 5) | 0.86 | 0.033 |
| | PreFecal (n = 5) | PostFecal (n = 9) | 0.32 | 0.029 |

**Table S3: Sequence and taxonomy of 178 oral bacterial ASVs identified from healthy human individuals involved in the HMP dataset.**

See separate Excel table

**Table S4: Oral bacterial ASV, taxonomy, and total abundance in human feces of MSKCC allo-HCT recipients. a**, Sequence and taxonomy of 127 oral bacterial ASVs. **b**, Total relative and absolute abundances of oral and gut bacteria in fecal samples. We determined the oral or gut bacterial load (absolute abundance) by multiplying the total bacterial load by the respective oral or gut bacterial fraction (relative abundance). Bacterial load unit is 16S copies per gram of feces.

See separate Excel table

**Table S5: Antibiotics associated with intestinal domination by oral bacterial ASVs in MSKCC allo-HCT recipients.** A total of 291 patients with at least 10 samples between day -10 and 40 relative to transplantation were included. The hazard ratios quantify the relative risk of intestinal domination by any single oral ASV that exceeds 30% in relative abundance compared to no domination. Oral and intravenous vancomycin were separated because vancomycin does not reach adequate levels in human gut when given intravenously. Distinct routes of administration were combined for all other antibiotics. *P* values were corrected for multiple comparisons using FDR. Significant associations ($P < 0.05$) are highlighted in red. CI: confidence interval.

| Antibiotic | Hazard ratio (95% CI) | *P* value |
|---|---|---|
| macrolide derivatives | 4.5e-8 (>0) | 0.996 |
| metronidazole | 0.26 (0.04-1.93) | 0.568 |
| fluoroquinolones | 0.45 (0.26-0.77) | 0.022 |
| sulfamethoxazole/trimethoprim | 0.52 (0.06-4.27) | 0.720 |
| aztreonam | 0.63 (0.15-2.68) | 0.720 |
| cephalosporins | 0.93 (0.50-1.72) | 0.880 |
| intravenous vancomycin | 0.94 (0.59-1.49) | 0.880 |
| carbapenems | 1.36 (0.72-2.57) | 0.580 |
| linezolid | 1.57 (0.69-3.58) | 0.580 |
| oral vancomycin | 1.77 (1.18-2.64) | 0.022 |
| piperacillin/tazobactam | 2.24 (1.37-3.65) | 0.015 |
| aminoglycosides | 2.95 (0.39-22.18) | 0.580 |

**Table S6. Regression slope between relative abundance of selected bacterial genera and total bacterial load in feces of MSKCC allo-HCT recipients.** Bacterial genera that dominated at least 100 fecal samples with a relative abundance exceeding 30% were selected for linear regression analysis in log-log space. In the table, we presented the mean relative abundance of each genus across HMP samples collected from the gastrointestinal tract, oral cavity, nasal cavity, skin, and urogenital tract. The five genera with the highest mean relative abundance in the gastrointestinal tract, compared to the other four body sites, are highlighted in red. Notably, they all exhibited positive slopes.

See separate Excel table

**Table S7: High-quality genomes of *Streptococcus* species assembled from shotgun metagenomics data.** Metagenome-assembled genomes (MAGs) were constructed from 22 fecal samples from MSKCC allo-HCT recipients. The iRep value is an index of replication and serves as an indicator of bacterial replication rate[2]. iRep values that did not meet all genome and mapping quality requirements are marked as “n/a”. Assembled MAGs containing *Streptococcus* ASV_8 are highlighted in red.

See separate Excel table

**Table S8: Oral bacterial fraction in feces is associated with survival of MSKCC allo-HCT recipients**. All-cause mortality and GVHD-related mortality were analyzed by a Cox proportional hazard model and a Fine-Gray subdistribution hazard model, respectively. Both models were adjusted for *Enterococcus* absolute abundance, age, underlying disease, graft source, and intensity of conditioning regime. *P* values were FDR-corrected for multiple comparisons. Abbreviations: CI (confidence interval), AML (Acute Myeloid Leukemia), ALL (Acute Lymphocytic Leukemia), MDS (Myelodysplastic syndrome), MPN (Myeloproliferative neoplasm).

| Endpoint event | Covariate | Hazard ratio | 95% CI | *P* value |
|---|---|---|---|---|
| All-cause mortality | Oral bacterial fraction | 4.02 | 2.11-7.68 | 6.3e-5 |
| | log(*Enterococcus* absolute abundance) | 1.19 | 1.13-1.25 | 3.5e-11 |
| | Age | 1.03 | 1.02-1.04 | 1.0e-12 |
| | Disease | | | |
| | AML/ALL/MDS/MPN | Reference category | | |
| | Others | 1.30 | 1.02-1.65 | 0.053 |
| | Graft source | | | |
| | T-cell depletion | Reference category | | |
| | Unmodified | 1.21 | 0.92-1.58 | 0.193 |
| | Cord | 1.33 | 0.89-1.99 | 0.193 |
| | Conditioning intensity | | | |
| | Ablative | Reference category | | |
| | Non-ablative | 0.46 | 0.30-0.72 | 0.001 |
| | Reduced intensity | 0.90 | 0.67-1.21 | 0.491 |
| GVHD-related mortality | Oral bacterial fraction | 4.23 | 1.69-10.6 | 0.006 |
| | log(*Enterococcus* absolute abundance) | 1.22 | 1.13-1.32 | 1.5e-6 |
| | Age | 1.03 | 1.01-1.04 | 2.3e-4 |
| | Disease | | | |
| | AML/ALL/MDS/MPN | Reference category | | |
| | Others | 1.31 | 0.90-1.91 | 0.319 |
| | Graft source | | | |
| | T-cell depletion | Reference category | | |
| | Unmodified | 1.07 | 0.67-1.70 | 0.777 |
| | Cord | 1.43 | 0.76-2.70 | 0.420 |
| | Conditioning intensity | | | |
| | Ablative | Reference category | | |
| | Non-ablative | 0.90 | 0.47-1.72 | 0.777 |
| | Reduced intensity | 1.14 | 0.71-1.84 | 0.777 |

**Table S9: Microbiome datasets used in this study.**

See separate Excel table

**Supplementary Note 1: Theoretical relationship between oral bacterial fraction and total bacterial load in fecal samples**

By definition, the relative and absolute abundance of oral and gut bacteria in a fecal sample are related through the following equation:

$$F_{total} = \frac{F_{oral}}{f_{oral}} = \frac{F_{gut}}{1 - f_{oral}} \qquad \text{Eq. S1}$$

Here, $f_{oral}$ represents the relative abundance of oral bacteria, $F_{oral}$ and $F_{gut}$ represent the absolute abundance of oral and gut bacteria respectively, and $F_{total}(= F_{oral} + F_{gut})$ is the total bacterial load.

<u>Pure *Marker* hypothesis</u>: When an increase in $f_{oral}$ is solely driven by gut bacterial depletion, $F_{oral}$ remains constant (let the constant be $K_1$). In this scenario, Eq. S1 can be rewritten as

$$F_{total} = \frac{K_1}{f_{oral}} \qquad \text{Eq. S2}$$

or on the log-log scale (base $b$)

$$log_b\, F_{total} = log_b\, K_1 - log_b\, f_{oral} \qquad \text{Eq. S3}$$

Eq. S3 indicates that the derivative of log-transformed total bacterial load with respect to log-transformed oral bacterial fraction is -1:

$$\frac{d\, log_b\, F_{total}}{d\, log_b\, f_{oral}} = -1 \quad (pure\ Marker\ hypothesis) \qquad \text{Eq. S4}$$

<u>Pure *Expansion* hypothesis</u>: When an increase in $f_{oral}$ is solely driven by absolute expansion of oral bacterial population, $F_{gut}$ remains constant (let the constant be $K_2$). In this scenario, Eq. S1 can be rewritten as

$$F_{total} = \frac{K_2}{1 - f_{oral}} \qquad \text{Eq. S5}$$

or on the log-log scale (base $b$)

$$log_b\, F_{total} = log_b\, K_2 - log_b(1 - f_{oral}) \qquad \text{Eq. S6}$$

From Eq. S6, we found that the derivative of log-transformed total bacterial load with respect to log-transformed oral bacterial fraction is positive:

$$\frac{d\, log_b\, F_{total}}{d\, log_b\, f_{oral}} = \frac{f_{oral}}{1 - f_{oral}} > 0 \quad (pure\ Expansion\ hypothesis) \qquad \text{Eq. S7}$$

**Supplementary Note 2: Validation of oral bacterial ASVs identified from healthy individuals in patients with inflammatory bowel disease**

Since human microbiota composition is body site-specific, we hypothesized that the reference set of 178 oral ASVs identified from a large cohort of healthy individuals recruited by the Human Microbiome Project (HMP) can be applied to other healthy individuals and patients for the detection of oral bacteria in their gut. To test this hypothesis, we analyzed a publicly available dataset from patients with inflammatory bowel disease (IBD) and their healthy controls[3]. This dataset includes paired fecal and saliva samples from a total of 43 healthy controls (HC), 16 patients with Crohn's disease (CD), and 42 patients with ulcerative colitis (UC).

By varying three cutoff parameters used to identify oral ASVs from HMP, we showed that the estimated oral fraction in the fecal samples of the IBD cohort participants is largely robust against variations in these parameter values (Extended Data Fig. 5a). Among the three cutoffs, the cutoff for mean relative abundance ($\theta_a$, see Methods for details) has the strongest impact. Across all healthy individuals, the mean oral bacterial fraction in feces was found to be 1.2%, which closely aligns with a previously reported estimate of 2%[4]. This fraction exhibited a nearly three-fold increase to 4.2% and 4.3% in CD and UC patients respectively, supporting the notion that IBD is associated with an enrichment of oral bacteria in the gut[5]. Most importantly, more than 84% (mean values: 92.2% for HC, 89.8% for CD, and 84.1% for UC) of the oral ASVs detected in the fecal samples from the reference set were also found in their corresponding saliva samples (Extended Data Fig. 5b). In addition, these oral ASVs accounted for over 87% (mean values: 93.4% for HC, 94.6% for CD, and 87.8% for UC) of the estimated total relative abundance of oral bacteria in the feces (Extended Data Fig. 5c). In summary, we validated that the reference set of oral ASVs identified from the HMP dataset can be used to infer oral ASVs in the fecal samples from other non-HMP healthy individuals and patients even in the absence of paired oral samples.

**Supplementary Note 3: Discussion on associations between biofilm-forming capacity of *Streptococcus*, *Actinomyces* and *Abiotrophia* and their fecal relative abundance in MSKCC allo-HCT recipients**

All three bacterial genera have the ability to form biofilm in the oral cavity[6,7]. However, it remains uncertain if they can form biofilms in the lower gastrointestinal tract. *Streptococcus thermophilus*, the most dominant *Streptococcus* species in the MSKCC allo-HCT cohort (see Table S7), is a poor biofilm producer due to its inability to firmly attach to surfaces[8]. Consistently, exopolysaccharides derived from *S. thermophilus* do not interact with mucin[9], indicating that this species may be unable to form biofilm in colonic mucus.

Furthermore, we used PICRUSt2[10] to predict KEGG[11] pathway abundances from 16S amplicon sequencing data (10,433 samples). The biofilm-forming potential of intestinal microbiota was quantified by the sum of relative abundance of three KEGG pathways containing the keyword "biofilm" (ko05111, ko02025, ko02026). We observed a positive, but statistically insignificant, Spearman correlation between this biofilm index and the relative abundance of oral *Streptococcus* (Spearman's rho = 0.004, $P = 0.661$). For oral *Actinomyces* and *Abiotrophia*, the correlations were statistically significant but negative (Spearman's rho = -0.075 and -0.056, $P$ = 9.1e-14 and 2.7e-8, respectively). Using a subset of 3,108 samples with 16S qPCR data, we found that the biofilm-forming capacity is positively associated with the total bacterial load (Spearman's rho = 0.073, $P$ = 6.3e-5). Here, all $P$ values have been adjusted for multiple comparisons.

Taken together, although we cannot entirely rule out the possibility, both literature studies and our findings do not provide evidence for a link between intestinal dominations by the three oral genera and their biofilm-forming capacity.

**Supplementary Note 4: Discussion on associations between antibiotic exposure and oral bacterial domination in the fecal samples from MSKCC allo-HCT recipients**

During allo-HCT, most patients received multiple antibiotics for prophylactic and treatment purposes for different periods. Due to different spectra of antibiotics, their administration led to diverse dynamics of oral bacteria translocated to the intestine, as observed in fecal samples by 16S rRNA amplicon sequencing (Fig. 3a). To isolate the individual effect of each antibiotic, we used a time-dependent Cox proportional hazards model (see Methods in the main text for details). The Cox model revealed that piperacillin-tazobactam, a combination of β-lactam and β-lactamase inhibitor with a broad spectrum of antibacterial activity, had the most significant positive influence on intestinal domination by any single oral ASV with a relative abundance that exceeds 30% (Table S5; Hazard ratio = 2.24; 95% confidence interval, 1.37-3.65; $P = 0.015$). This association aligns with a previous finding showing that piperacillin-tazobactam caused the most pronounced depletion of anaerobic gut commensals in the same patient cohort[12].

To validate the anaerobe-depleting effect of piperacillin-tazobactam, we reanalyzed a previous study[1] that investigated the dynamics of gut microbiota during and after total gut decontamination (using oral piperacillin-tazobactam) and selective gut decontamination (using oral polymyxin-neomycin) in children undergoing allo-HCT at the Leiden University Medical Center in the Netherlands. Indeed, the children who received piperacillin-tazobactam exhibited significantly higher total fraction of oral bacteria in their feces compared to those who received oral polymyxin-neomycin across all transplantation stages (Fig. S2).

Our analysis also revealed a significant negative association between fluoroquinolone antibiotics with oral bacterial domination (Table S5; Hazard ratio = 0.45; 95% confidence interval, 0.26-0.77; $P = 0.022$). Fluoroquinolones are commonly used as prophylactic agents to reduce the incidence of gram-negative bacterial infections, including *Pseudomonas* and *Enterobacteriaceae*, in patients with neutropenia[13]. One possible explanation for this negative association is that fluoroquinolones are generally more effective against aerobic and facultative anaerobic bacteria, such as those found in the oral flora, than against anaerobic bacteria[14]. Another potential explanation is that all allo-HCT recipients were initially prescribed fluoroquinolone prophylaxis but may be switched to other agents, such as piperacillin-tazobactam, when fevers or bloodstream infections occur. As a result, fecal samples exposed only to fluoroquinolones may be enriched for samples collected before acute febrile episodes or from patients with more benign treatment courses who never got a fever and thus remained on fluoroquinolone prophylaxis.

### Supplementary Note 5: Slow growth of *Streptococcus* ASV_8 in fecal samples from MSKCC allo-HCT recipients

Among all identified oral ASVs, *Streptococcus* ASV_8 had the highest mean relative abundance among all fecal samples of the MSKCC allo-HCT recipients. We previously published shotgun metagenomics data[15] for 395 fecal samples from these patients. Among the 395 samples, 19 contains at least 10% *Streptococcus* ASV_8, as indicated by the paired 16S rRNA sequencing. Using a published bioinformatic pipeline (see Methods in the main text for details), we obtained 22 high-quality metagenome-assembled genomes (MAGs) of *Streptococcus spp.* from these 19 metagenomic samples. Among the 22 MAGs, four contains ASV_8, and all were annotated as *S. thermophilus*. We then calculated the replicate rate of these MAGs through iRep[2]. The iRep index can accurately estimate the ratio between the coverage at the origin and terminus of replication, which is proportional to replication rate. We found an averaged iRep index of 1.35 (Table S7), indicating that, on average, only 35% cells are replicating. This iRep-based estimation suggests that bacteria containing *Streptococcus* ASV_8 grew slowly in the intestine of MSKCC allo-HCT recipients.

**Supplementary Note 6: Oral bacterial fraction in feces predicted an increased risk of patient mortality after allo-HCT**

Previous analysis of the MSKCC allo-HCT cohort has established a link between intestinal expansion of *Enterococcus* and higher patient mortality[16]. In this study, we aimed to investigate whether the oral bacterial fraction in feces is also associated with mortality for 1,268 allo-HCT recipients with available survival information. We employed a Cox proportional hazard model adjusted for confounders that include *Enterococcus* absolute abundance, age, underlying diseases, graft source, and conditioning regimen (see Methods in the main text for details). The Cox model revealed that a higher oral bacterial fraction was associated with a higher risk of all-cause mortality following allo-HCT (Table S8; Hazard ratio = 4.02; 95% confidence interval, 2.11-7.68; $P$ = 6.3e-5). Since graft-versus-host disease (GVHD) is a major complication after allo-HCT, we further explored whether GVHD contributed to the observed association between oral bacterial fraction and all-cause mortality. Among 462 patients who died within two years after allo-HCT, 168 developed GVHD. Using a Fine-Gray competing risk regression model (see Methods in the main text for details), we further determined that the total fraction of oral bacteria predicted an elevated risk of GVHD-related mortality within two years (Table S8; Hazard ratio = 4.23; 95% confidence interval, 1.69-10.6; $P$ = 0.006). These associations with patient survival demonstrate that oral bacterial fraction in feces serves as a quantitative measure of microbiome damage that adversely affects host health.
